# Development of a network formation assay for developmental neurotoxicity hazard screening using 3D human iPSC derived BrainSpheres

**DOI:** 10.1101/2025.06.03.657699

**Authors:** SL Boyd, Mayil Vahanan M, J Monroe, JA Conley, KE Carstens, TJ Shafer

**Affiliations:** Center for Computational Toxicology and Exposure, USA Environmental Protection Agency, Research Triangle Park, NC; Oak Ridge Institute for Science and Education, Oak Ridge, TN

## Abstract

Exposure of the developing brain to environmental neurotoxicants can result in permanent alterations in structure and/or function. To investigate the effects of chemical exposures on neurodevelopment, the human induced-pluripotent stem cell (iPSC)-derived neural BrainSphere model has been utilized due to its ability to form mature neuronal populations and exhibit spontaneous electrical activity. To model network formation for developmental neurotoxicity screening, developing BrainSpheres were plated on high-density microelectrode arrays (hdMEA) three weeks after beginning differentiation. Starting two days post-plating, BrainSpheres were treated three times per week with compounds known to disrupt *in vitro* network formation (i.e. assay positive controls; loperamide, dieldrin and deltamethrin), or with an assay negative control, glyphosate, expected to have no effect. For 29 days, BrainSphere activity was recorded to measure neural network activity, general activity, and features of action potential propagation. Concentration-dependent disruption in neural network formation was observed for positive controls at concentrations below cytotoxicity. Dieldrin, deltamethrin, and loperamide exposure disrupted several features of general activity, neural network formation, and action potential propagation. BrainSpheres on hdMEAs detected chemically induced perturbations in neural network formation and may represent a valuable complex *in vitro* model useful for developmental neurotoxicity screening.

## Introduction

Neurodevelopment entails a complex series of processes that give rise to multiple different neural cell populations, each with specific functions contributing to neural circuits and behavior. Some of these neurodevelopmental processes include proliferation, migration, differentiation, synaptogenesis, and the formation of neural networks. Each of these processes is highly regulated and perturbations, such as those induced by a neurotoxicant, can result in molecular, structural or behavior changes. Since the developing brain is particularly vulnerable, such disruptions can contribute to the characteristic symptoms exhibited in neurodevelopmental disorders (Rice and Barone, 2000; Giordano and Costa, 2012). Developmental neurotoxicity (DNT) has become a concern as neurodevelopmental disorders are becoming more prevalent worldwide and some epidemiological studies have reported associations between environmental exposures and neurodevelopmental disease (Chakrabarti and Fomebonne 2005; McDonald and Paul 2010; U.S. Environmental Protection Agency, 2024a). Due to the complexities in neurodevelopmental processes, there are several challenges to assessing DNT hazard including the inability to completely model the intricacies of the human brain, difficulty isolating the appropriate exposure window, inter-species differences, difficulty extrapolating between *in vivo* and *in vitro* models, as well as many others. Traditionally, DNT hazard assessment uses *in vivo* guideline studies (U.S. Environmental Protection Agency, 1998; Organisation for Economic Co-operation and Development, 2007). To date, of the thousands of chemicals to which humans are exposed, only approximately 230 have been evaluated in DNT guideline studies thus far (Crofton and Mundy, 2025). In addition, these studies are resource intensive, require high numbers of animals, and results are often difficult to interpret and extrapolate to humans, as they lack any indication of the underlying biological changes contributing to alterations in the endpoints evaluated within them. (Tsuji and Crofton, 2012).

The need for chemical safety information on hundreds to thousands of chemicals, combined with the goal to improve upon the understanding of biological alterations underlying and human relevancy of toxicological testing approaches, particularly for DNT, has driven the development of new approach methodologies (NAMs) (Environmental Protection Agency, 2024b). NAMs are assays and technologies that can be used for toxicology assessment aimed at improving human relevancy, thereby reducing the usage of (or reliance on) animal models. Recently, The Organisation for Economic Co-operation and Development (OECD) recognized a DNT *in vitro* battery (DNT-IVB) that comprises 17 unique assays (Organisation for Economic Co-operation and Development, 2017; 2023). These assays evaluate key neurodevelopmental processes, and the collective information gained from all assays provides useful information regarding DNT hazard (Sachana et al., 2021). The DNT-IVB or specific assays in the DNT-IVB have been used to screen ∼500 compounds for screening, prioritization, and contributions to weight-of-evidence (Carstens et al., 2023; Kreutz et al, 2024; Dobreniecki et al., 2022), and several scientific advisory panels and expert groups have concluded these applications of the DNT-IVB are appropriate (Juberg et al., 2023; U.S. Environmental Protection Agency, 2020; The Organisation for Economic Co-operation and Development, 2023). Additionally, these reviews were also consistent in noting that DNT NAMs should evolve with the science, and that future improvements to the battery should include increasing human relevancy, biological complexity, as well as the incorporation of a tiered testing approach.

The neural network formation assay (NFA) is unique to the DNT*-*IVB since it is currently the only assay that evaluates chemical perturbations on neural function rather than morphological or phenotypic assessments (Brown et al., 2016; Frank et al., 2017). The NFA uses primary rat cortical neurons grown on microelectrode arrays (MEAs) containing 16 electrodes per well that measure spontaneous electrical activity of the cells (Brown et al., 2016). Within the first week after plating, the neural networks that form exhibit low rates of individual action potential firing that are uncoordinated, and activity is observed only a few of the 16 electrodes in each well. However, as the networks mature over the course of 12 days *in vitro*, the neurons across all electrodes exhibit increased firing rates, bursts of action potentials on individual electrodes, and synchronous bursts of activity across all electrodes (network bursts) indicating an interconnected network (Cotterill et al., 2016). Network bursts are the biological underpinnings of cognitive processes that include learning and memory (Buschman and Kastner, 2015; Korte and Schmiz, 2016; Salinas and Sejnowski, 2001). Characterization of the NFA has revealed expression of glutamatergic, GABAergic, and astrocytic populations in the cortical culture (Brown et al., 2016; Frank et al., 2017). Although the mechanisms measured in the NFA are analogous to human cognitive development, the established model has noted areas for improvement. The NFA is a two-dimensional monolayer culture system that uses primary rodent cortical cells within our implementation. Therefore, a goal of this work was to improve upon the human relevance and biological complexity of the NFA by evaluating a methodological change to utilize human iPSC-derived BrainSpheres as an innovative compliment for two-dimensional cultures of rodent cells.

BrainSpheres are made from iPSC-derived cells that have been formed into 3-dimensional (3D) structures. The 3D structure improves upon the cytoarchitecture of traditional monolayer cultures as they spontaneously self-assemble, forming structures with inter-connected complex neural networks (Pamies et al., 2016; Hartmann et al., 2023; Lislien et al., 2025). Furthermore, the iPSC-derived BrainSpheres incorporate a heterogenous cell population including RNA expression of stem cell and proliferating cells early on during development; eventually these markers diminish as the BrainSpheres mature (Pamies et al., 2017). BrainSpheres also express markers of astrocytes and oligodendrocytes, as well as neuronal subtypes including glutamatergic, GABAergic, and dopaminergic cells (Pamies et al., 2017). Additional work has been conducted to quantify relative gene expression levels for neuronal subtype markers, ion channels, and neurotransmitter receptors and transporters confirming this heterogeneity and complexity of the model (Hartmann et al., 2023). Furthermore, Hartmann and colleagues demonstrated and characterized that BrainSpheres are functionally active and can develop spontaneous electrical activity as well as network bursts (Hartmann et al., 2023).

A recent publication by Carstens and colleagues demonstrated that BrainSpheres and their ability to produce spontaneous electrical activity can be leveraged to assess perturbations in the context of neurotoxicity (Carstens et al., 2025). More specifically, Carstens and colleagues plated multiple BrainSphere (at seven weeks post induction of differentiation (PID)) into one well of an hdMEAs and evaluated perturbations to general and network activity, representing a more mature brain model of neurotoxicity. Although these results ensure that the BrainSphere model can be utilized for assessing neurotoxicity, the current study is specifically focused on developmental neurotoxicity. In early stages of brain development, spontaneous electrical activity levels are high as the brain is developing synaptic connections, and as the brain matures the activity declines (Banies and Landgraf, 2021; Levitt, 2003, Tau et al., 2009). Therefore, the current study aims at modeling an early phase of neurodevelopment when spontaneous electrical activity is peaking to ensure that the exposure paradigm recapitulates a critical phase in neurodevelopment in which the developing brain is most vulnerable to toxicological perturbations (Rock and Patisaul, 2019).

The current study sought to explore whether BrainSpheres could be developed into a suitable assay for DNT screening, potentially as a higher tiered test to provide a more in-depth characterization of toxicological hits from the DNT-IVB. The recommendations of Crofton et al., 2011 were followed to develop the assay. BrainSpheres were treated with a small set of assay positive controls and an assay negative control as the BrainSpheres formed neural networks on MEAs. These control compounds were selected because they previously have been shown to alter network formation in rat primary cells grown on MEAs. Starting at two-days post-plating, BrainSpheres were treated with three assay positive controls (loperamide, dieldrin, deltamethrin) or one assay negative control (glyphosate), which were selected due to their well characterized effects (or lack of effects) in *in vitro* NFA studies using primary rat cortical neurons (Brown et al., 2016; Frank et al., 2017; Shafer et al., 2019). Loperamide is a µ-type opioid receptor antagonist, while dieldrin and deltamethrin are organochlorine and pyrethroid ester pesticides, respectively. Glyphosate is an organophosphorus herbicide that has been tested in the NFA multiple times without any activity, making it a suitable negative control for this assay (Carstens et al., 2022). Furthermore, glyphosate has been tested in other assays in the DNT-IVB and has been shown to be a reliably negative compound in DNT assays based on assay performance and the current literature landscape (Martin et al., 2022).

BrainSpheres were differentiated for three weeks, then plated on high density microelectrode arrays (hdMEAs) for up to 29 days. Throughout the 29 days on the hdMEA, the BrainSpheres were exposed to a chemical treatment and electrical activity in the BrainSpheres was recorded to measure network formation, general activity, and features of action potential propagation and axonal morphology. In comparison to the standard MEAs which contain 16 electrodes per well, the hdMEAs have 26,400 electrodes per well, greatly increasing both the spatial and temporal resolution. This method therefore leverages biologically complex, human cell-derived 3D model and hdMEA for the purposes of evaluating an assay to assess neural network formation with potential use in a tiered testing strategy for DNT. Therefore, we hypothesize that chemical exposure in the BrainSphere/hdMEA assay will result in disruptions to network formation, paralleling results previously established in the NFA, and the assay will provide additional insight features such as action potential propagation.

## Methods

### Cell Culture and Maintenance

Induced Pluripotent stem cells (iPSCs) were originally generated from the following cell lines CCD1079Sk (ATCC, CRL2097™), IPS IMR90 (WiCELL), and ATCC-DYP0730 human Induced Pluripotent Stem (IPS) cells (ATCC, ACS1003™). Generation of iPSCs, Neural Progenitor Cells (NPCs) production, and BrainSphere differentiation were completed as previously described (Pamies et al., 2017; Wen et al., 2014). Briefly, iPSCs were treated with neural induction supplement (Gibco, A1647801) starting on day 2 post-plating and maintained for seven days forming NPCs. NPCs were expanded and cryopreserved at generations four through eight. After at least two generations, NPCs were seeded at a density of 2×10^6^ cells/well on ultra-low attachment 6-well culture plates in expansion media (49.5% Advanced DMEM/F-12 (Gibco, 12634), 49.5% Neurobasal Medium (Gibco, 21103049), 1% Neural Induction Supplement (Gibco, A1647801)) and placed on an orbital shaker at 88rpm in a 5% CO_2_ humidified atmosphere at 37°C, allowing BrainSphere self-assembly. Two days post-initiation of spheroid formation, BrainSphere differentiation was initiated by replacing the medium with differentiation medium containing Neurobasal Plus medium (Gibco, A35829) supplemented with 5% B-27 Plus supplement (Gibco, A35828), 1% GlutaMAX^TM^ (Gibco, 35050061), 0.02 μg/ml human recombinant GDNF (Gemini, 300-121P), and 0.02 μg/ml human recombinant BDNF (Gemini, 300-104P). BrainSphere cultures were maintained for 3 weeks by conducting half-medium changes with differentiation medium twice a week prior to plating on hdMEAs at 5% CO_2_ and 37°C. While BrainSpheres were cultured in suspension, half-medium changes were completed twice a week. In parallel, additional 6-well plates containing free floating BrainSpheres were maintained for up to ten weeks to sample BrainSpheres for immunocytochemistry characterization (See Immunocytochemistry and confocal microscopy methods section below).

### hdMEA plating

At three weeks PID, Brain Spheres were plated on either 6-well or 24-well hdMEA plates (Maxwell BioSystems) (Comparison of the ontogeny of spontaneous electrical activity for BrainSpheres grown on 6 vs 24-well plates is shown in Supplemental Figure 1). Five days prior to plating, differentiation medium was added to each well of the hdMEA plates and stored in an incubator at 37 C, 5% CO_2_ for 48 hours (Figure 1). Next, medium was aspirated from each well and washed three times with sterile water.

**Figure 1.**
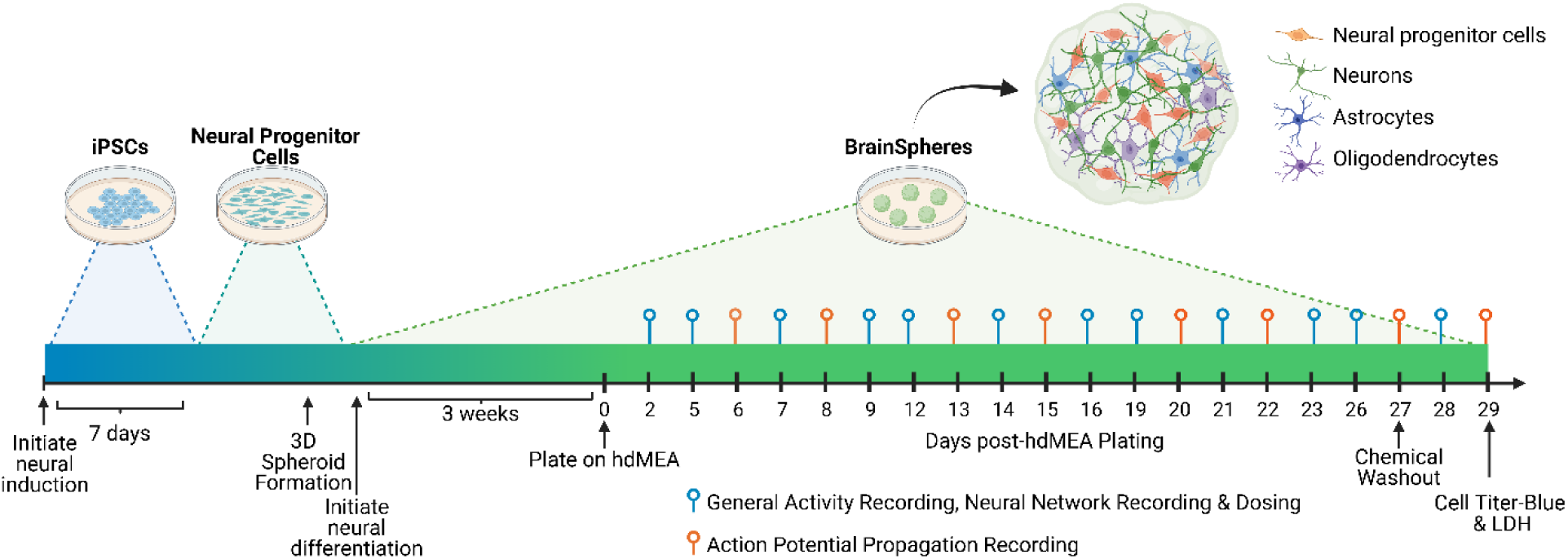
Experimental Design. Pluripotent stem cells are induced for 7 days to form neural progenitor cells. Neural progenitor cells are formed into 3D spheroids, and two days later neural differentiation is initiated to form BrainSpheres. BrainSpheres develop a heterogenous cell population containing neural progenitor cells, neurons, astrocytes, and oligodendrocytes. BrainSpheres are maintained for 3 weeks, then plated onto high density microelectrode arrays for 29 days. Three times a week, BrainSpheres are recorded using activity and network recordings, and dosed with chemicals during a complete medium change. Two times a week, BrainSpheres are recorded using axon tracking software to analyze characteristics of action potential propagation. On Days on Plate (DOP) 27, medium is aspirated and replaced with fresh medium as a chemical washout. One set of general activity, neural network, and action potential propagation recordings are completed on the washed-out BrainSpheres. On DOP 29, BrainSpheres are assessed for cytotoxicity using CellTiter-Blue® and LDH assays.

Plates were next coated with 0.07% Poly(ethyleneimine) solution (PEI, Sigma-Aldrich, 181978) in 1x borate buffer (Thermo Scientific, 28341) and incubated overnight. PEI was then aspirated from each well and wells were rinsed once with water. Following, 70µl of 0.02mg/ml laminin (Sigma-Aldrich, L2020) was diluted in sterile DNAse/RNAse free water (Invitrogen, 10977015) was added to the center of each well and incubated for one hour.

On the day of plating, hdMEA plates were set on a warming plate set to approximately 37 °C and differentiation medium (500µl for 24-well plates and 1ml for 6-well plates) was added to each well. One BrainSphere was pipetted into each well and gently manipulated to the center of each well using a wide-bore, 200µl pipette tip. After plating BrainSpheres on the hdMEA plates, the plates were maintained on the warming plate for approximately four hours to allow attachment. During the four-hour incubation, a lid was placed over the plates allowing 5% CO_2_ to flow over each hdMEA plate. Once the BrainSpheres were attached, the plates were sealed with BreatheEasy sealing membranes (Sigma-Aldrich, Z380059) and returned to the incubator. BrainSpheres were plated on a combination of both 6-well and 24-well hdMEA plate types and all data were normalized to the BrainSphere’s respective plate type (either 6-well or 24-well).

Prior to each recording, BrainSpheres that were plated on hdMEAs were visually inspected to ensure that each BrainSphere was deemed acceptable for experimentation. The acceptability criteria ensured that each BrainSphere was centered on the hdMEA chip, looked round in appearance, appeared relatively uniform in size, was properly attached (not free floating) and lacked debris in the well. If a BrainSphere did not meet these acceptability criteria at any point in the experiment, the well was removed from all analysis.

### Chemical Treatment and Media Exchanges

Starting on day-on-plate (DOP) 2, BrainSpheres were treated with chemicals three times a week during full media changes until DOP 26 without a wash step, which occurred immediately following hdMEA recordings (Figure 1). For each chemical treatment, three to four biological replicates were tested.

Biological replicates were defined as one individual BrainSphere or one individual well of the hdMEAs (BrainSpheres were plated one per well) each coming from a separate NPC differentiation and spheroid formation batch. BrainSpheres were plated on a combination of 24-and 6-well hdMEA plates, due to availability of plates at the time of experimentation.

For 24-well plates, 4 chemicals (5 concentrations and one vehicle control) were tested per plate (See Supplemental Figure 2 for example plate layouts). Each chemical was tested in a different row and each concentration was tested in a different column. The vehicle controls (either DMSO or water) were position in column 2 to avoid any edge effects in control wells. The lowest concentration was tested in column 1. The remaining concentrations were tested in columns 3-6 at increasing concentrations. For 6-well plates, only one chemical (5 concentrations and one vehicle control) was tested per plate. The vehicle control was tested in well A2 to avoid potential edge effects. The lowest concentration was tested in A1, and the remaining concentrations were tested in wells A3, B1-B3 at increasing concentrations. Each chemical’s replicates were tested in a combination of 6-well and 24-well plates.

Stock solutions of dieldrin (Chem Service, N-11688, cas: 60-57-5), deltamethrin (Chem Service, N-11579, cas: 52918-63-5), and glyphosate (Chem Service, N-12133, cas: 1071-83-6) were aliquoted and stored at 100mM concentrations in dimethylsulfoxide (DMSO) or in water. Loperamide (Sigma-Aldrich, 1370000, cas:34552-83-5) was stored at 3mM in DMSO. To prepare for dosing, a 96-well plate (ThermoFisher, 2305-11) was utilized as the “dilution plate” and a deep-well plate (ThermoFisher, 278606) was utilized as the “dosing plate.” In the dilution plate, chemicals were serially diluted in either DMSO (dieldrin, deltamethrin, and loperamide) or water (glyphosate only) to form concentrations that are 1,000x the final dilution. Next, 495µl of differentiation medium was added to each well of the dosing plate and 5µl from the dilution plate was added and mixed well to achieve a concentration 10x the desired final chemical concentration. The final DMSO or water concentration was 0.1%.

After hdMEA recordings, differentiation medium from each well was gently aspirated and replaced with 450µl for 24-well plates or 900µl for 6-well plates of differentiation medium. To dose, 50 or 100µl was pipetted from the dosing plate into the 24-well or 6-well hdMEA plate, respectively, for a final chemical concentration of 1x. Plates were sealed with BreatheEasy plate sealers and returned to the incubator.

### hdMEA Recording

Three times a week starting on DOP2, spontaneous electrical activity of the BrainSpheres was evaluated with the Activity and Network recording modules in the MaxLab Live (version 24.1.2) using the MaxTwo hdMEA platform (Maxwell Biosystems, Zurich, Switzerland)(Figure 1). All hdMEA recordings were conducted with a high pass filter set to 1Hz, a gain of 512x, spike threshold of 5.0, number of filter taps set to 100, Beta of 5.0, highpass cutoff of 300, and a lowpass cutoff of 3,000. Activity recordings were first completed by scanning every other electrode across the entire array for a total of 15 minutes to measure parameters of general activity such as active electrode area or firing rate (Figure 2A). Analysis of activity recording data was set to the following parameters in MaxLab Live: firing rate threshold set at 0.1Hz, spike amplitude threshold at 20.00µV, and an Inter-Spike-Interval (ISI) set to 200ms. The Network recording module was then utilized to assess parameters of network bursts and clusters of network bursts (Figure 2B). In the network recording module, data was only collected from those electrodes that were identified as “active” in the activity recording module and were recorded for a total of 15 minutes.

**Figure 2.**
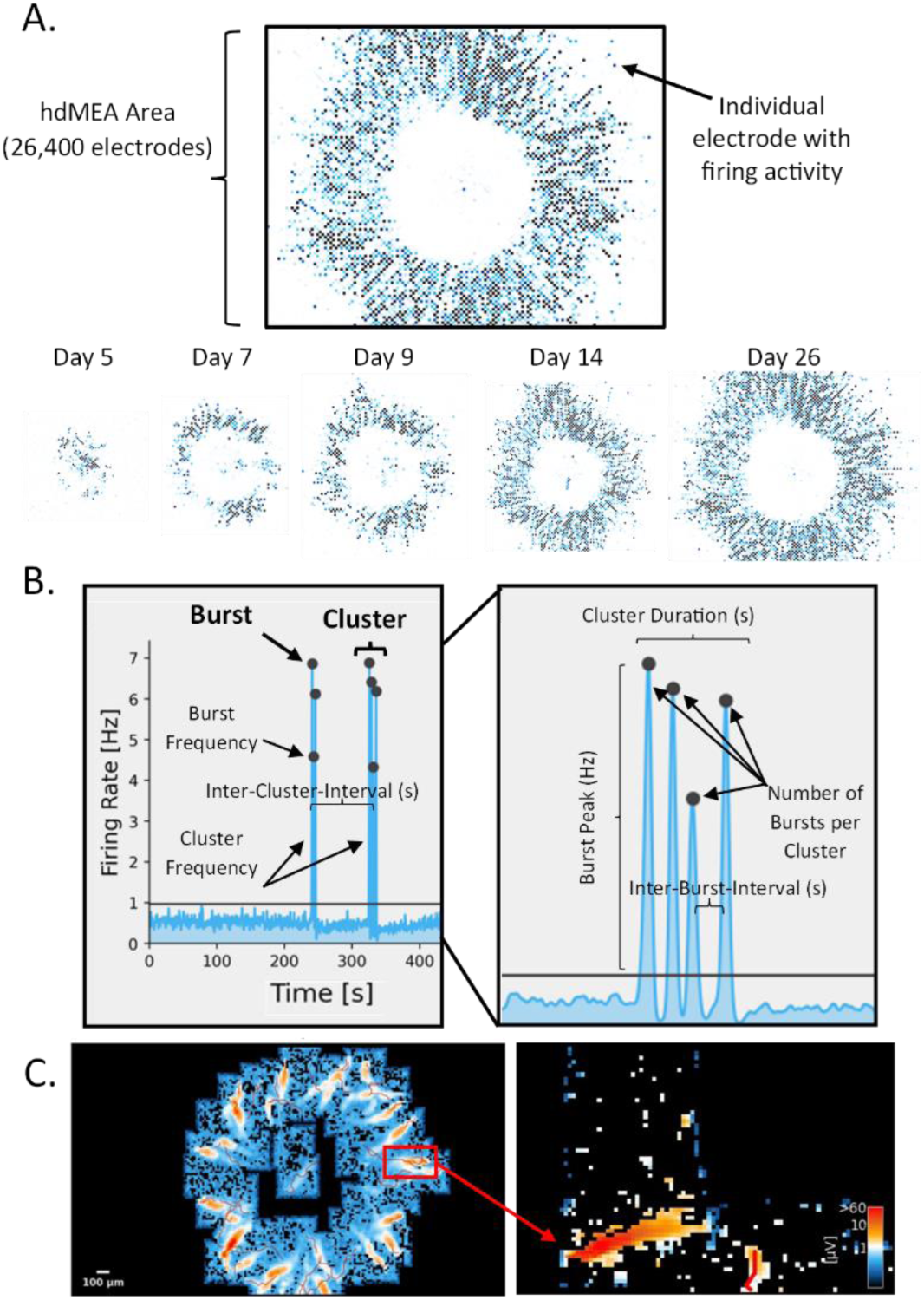
Overview of the three recording types for BrainSpheres plated on hdMEAs. A) A representative image of a BrainSphere plated on an hdMEA. The black rectangle is representative of the area of the hdMEA surface, and each point represents an electrode with firing activity. The bottom image of part A includes a time course of images of the firing activity of a BrainSphere on an hdMEA. B) A representative graph of firing rate (Hz) vs time for a BrainSphere plated on an hdMEA defining burst versus clustering activity. C) A representative image of axons detected in the MaxLab Live software.

Twice weekly, the Axon Tracking (action potential propagation) module was used to assess both functional and morphological features of action potential propagation (Figure 2C). For action potential propagation analysis, regions of interest were identified based on the last activity recording and axons were detected by setting the time window before an action potential to 1.5ms, the time after set to 5.0ms, the number of spikes threshold to 10, the footprint completeness threshold to 0.6, a latency threshold set of 0.0ms, and the radius of pixels set to 3.0.

### Coefficient of Variation Calculations

The coefficient of variation (CV) of controls for each endpoint was calculated by first normalizing by plate type (24 versus 6-well hdMEA plate), DOP, and endpoint. Normalized data was then transformed using trapezoidal integration across DOPs producing an AUC value. The coefficient of variation (CV) for each endpoint was computed using the normalized AUC values with the following equation:

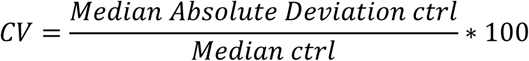

A CV value of 40% was determined as an appropriate threshold for investigating if an endpoint in the assay had low (CV < 40%) or high (CV > 40%) variability.

### Cytotoxicity Assays

On DOP 27, all media were aspirated from each well and washed with 500µl of fresh differentiation medium twice and replaced with 500µl of fresh differentiation medium for the remainder of the experiment to serve as a chemical washout. After the final recording, CellTiter-Blue® (Promega, G9090) and lactate dehydrogenase (LDH) (Promega, G1781) assays were conducted for each well (one BrainSphere/well). For the CellTiter-Blue® assay, the CellTiter-Blue® solution was first diluted 1:5 in differentiation medium. In the hdMEA plate, the medium was aspirated from each well and replaced with 100µl of the previously made CellTiter-Blue® medium solution. To prepare the CellTiter-Blue® assay plate, 150µl of CellTiter-Blue® medium solution was added to three wells as a blank in a white 96-well assay plate. Both the assay and hdMEA plates were then incubated for 1 hour. Following, 70µl was pipetted out of each hdMEA plate well and into the assay plate. The plate was read using a Fluorstar Optima (BMG LabTech) at a gain of 1,380x, excitation of 544nm, and emission at 590nm resulting in raw fluorescent values. Description of the analysis is listed in the data analysis and statistics methods section below.

Following the CellTiter-Blue® assay, the LDH assay was next performed in the same hdMEA plate. Since the half-life of LDH in medium is only about 9 hours, extracellular LDH content is not a reliable metric of cell death over a long period of time. Therefore, LDH in the remaining cell pellet was determined to provide a metric of the number of cells that remain following treatment, relative to control wells. To prepare the LDH plate, 500µl of lysis solution was mixed with 25 ml of Dulbecco’s 1X Phosphate Buffered Saline (DPBS) (Gibco, 14190-144) to make the lysis solution. As a blank control in the assay, 25µl of the lysis solution was added to three wells in a 96-well clear bottom plate. Two LDH positive controls were also added to the plate. The LDH positive control stock was diluted 1:250 and 1:2,250. For a neutral control, 50µl of differentiation medium was pipetted into three separate wells of the plate. In the hdMEA plate, the remainder of the CellTiter-Blue® solution was aspirated from each well. Next, 150µl of the lysis solution was added and mixed well, and then the plate was placed in the incubator for 30 minutes. Following, 50µl of the solution was pipetted out of the hdMEA plate and into the assay plate with an additional 50µl of reconstituted substrate and mixed well. Next, the plate was incubated for 30-minutes in the dark. At the end of this incubation period, 50µl of the stop solution was added to each well in the assay plate and the absorbance was measured using a VersaMax microplate reader (Molecular Devices) at a wavelength of 590nm. Description of the analysis is listed in the data analysis and statistics methods section below.

### Immunocytochemistry and Confocal Microscopy

BrainSpheres were fixed at weeks 3 and 7 PID by submersion in 4% paraformaldehyde (Thermo Scientific, 043368.9M, diluted in DPBS) for ∼12 hours then rinsed with DPBS at weeks three and seven. BrainSpheres were moved to 74µm, polyester membrane netwell inserts for the remainder of the staining process (Corning, 3477). Netwells containing BrainSpheres were washed three times with DPBS. Following these rinses, all DPBS was aspirated, replaced with 500µl of permeabilization solution (0.2% Triton-x (Alfa Aesar, A16046), 2% Bovine Serum Albumin (BSA) (Sigma-Aldrich, A7906, in DPBS), and BrainSpheres incubated at room temperature for 30 minutes. Permeabilization solution was next aspirated, BrainSpheres were rinsed with immunocytochemistry staining buffer (ISB) (.5% BSA, 0.05% sodium azide (Sigma-Aldrich, S2002), 0.01% saponin (Fluka Analytical, 47036) in DPBS) two times and then ISB was replaced with primary antibodies (Mouse-IgG1-β-III-Tubulin (Promega, G712A; 1:1,000) and Rabbit-IgG-PAX6 (Invitrogen, 42-6600; 1:200)) diluted in ISB and incubated for 24 hours at 4 °C. Following, primary antibody solution was aspirated, wells containing BrainSpheres were washed three times with ISB and ISB was replaced with a secondary antibody solution containing secondary antibodies diluted 1:500 in ISB (Goat-anti-Rabbit-IgG (H+L) 647 (Invitrogen, A21245), Goat-anti-Mouse-IgG (H+L) 488 (Invitrogen, A11029)) and incubated at room temperature for two-hours protected from light.

Following secondary incubation, BrainSpheres were rinsed with ISB twice, then ISB was replaced with ISB containing Hoechst (1:10,000) for ten minutes, rinsed once with ISB, and stored in DPBS. BrainSpheres were then transferred to a black, 96-well imaging plate (CellCarrier-96 ultra, 6055302, with 150µl of DPBS per well) and imaged with a confocal microscope (Nikon A1 Confocal) at 10x at the following wavelengths, 402nm,488nm, and 647nm.

### hdMEA Data Preprocessing

The overall data landscape included four chemicals tested, with five tested concentrations per chemical (plus vehicle controls), and a total of 32 unique endpoints from the three types of hdMEA recordings.

HdMEA raw data was exported from the Maxwell Software as csv files and imported into R (version 4.4.1). Briefly, raw csv files were exported from MaxLab Live (version 24.1.2) which contained all endpoints relating to general activity and action potential propagation. For the neural network endpoints, the number of spikes per burst, inter-burst-interval, and burst peak were provided by MaxLab Live software. However, a specific neural network pattern was observed in BrainSpheres resulting in a clustering of bursts (see Figure 2B). Since the parameters provided by the MaxLab Live software did not capture this clustering-like behavior, the remaining neural network level endpoints were manually derived using R statistical computing language. To analyze neural network level activity and manually derived additional endpoints, bursting activity was organized into clusters (Figure 2B).

Clusters were defined as grouping of bursts that occurred within an inter-burst-interval less than 7.5 seconds. Cluster-specific and burst-specific methods were then derived to characterize the activity in BrainSpheres. Endpoints for each type of recording and their descriptions are listed in table 1.

**Table 1.** High-Density hdMEA Endpoint Description (MaxWell BioSystems)

| Recording Type | Endpoint | Level of Data | Units | Brief Description |
| --- | --- | --- | --- | --- |
| **General Activity** | Active Area | Well | NA | The number of electrodes that have a firing rate larger than 0.1 Hz and a spike amplitude greater than 20 μV. |
|  | Inter-Spike-Interval |  | ms | The time between spikes. |
| | Spike Amplitude | | μV | The 90th percentile of the amplitude distribution for all detected spikes on that electrode. |
|  | Firing Rate |  | Hz | The number of spikes divided by the recording time. |
| **Neural Network** | Number of Spikes per Burst | Burst | NA | The number of spikes that fall between the beginning and end of a single burst on each electrode. |
|  | Inter-Burst-Interval |  | ms | The time between network bursts. |
|  | Burst Peak |  | Hz | The smoothed peak firing rate or the number of spikes over time, of the detected network burst. |
|  | Burst Frequency |  | Counts/min | The number of detected network bursts per minute. |
|  | Burst Duration |  | ms | The time between the beginning and end of a network burst. |
|  | Number of Bursts per Cluster | Cluster | NA | The number of network bursts per BrainSphere cluster. |
|  | Cluster Frequency |  | Counts/min | The number of clusters per minute. |
|  | Inter-Cluster-Interval |  | ms | The time between clusters. |
|  | Cluster Duration |  | ms | The time between the beginning and end of a cluster. |
| **Action Potential Propagation** | Branch Length | Branch | μm | Length of each axonal branch. The length is the collective distance between branch positions detected. |
|  | Conduction Velocity |  | m/s | The action potential propagation along the branch is velocity. Velocity is calculated from the linear regression fit of the collective distance between detected branch points and the corresponding spike negative peak times. |
|  | Number of Spikes per configuration |  | NA | The mean number of spikes for each recording configuration. Larger number represents a lower signal-to-noise proportion of the reconstructed signal, and therefore a greater footprint quality. |
|  | Firing Rate |  | Hz | The number of spikes divided by the recording time in a branch. |
|  | Conduction velocity coefficient of determination |  | NA | The R-squared value to evaluate the conduction velocity fit. |
|  | Longest latency from initiation site |  | ms | The amount of time for an action potential to propagate from the initiation site to the beginning of the branch. |
| | Longest distance from initiation site | | μm | The Euclidean distance from the starting point of the action potential initiation site to the beginning of the branch. |
|  | Fisher projection distance |  | NA | A metric of the separability between the main grouping of spikes and other spike groupings. |
|  | Silhouette Score |  | NA | A metric indicative of spike sorting quality. The score ranges from 0 to 1 and indicates the range between the main and remaining clusters. 1 represents clusters that are distinguishable. |
|  | Total Number of Spikes |  | NA | The total number of spikes in a cluster. |
|  | Completeness Score |  | NA | Indicative of the fraction of configurations containing greater than 10 spikes for computing a footprint. |
|  | Firing Rate | Neuron | Hz | The number of spikes divided by the recording time in a neuron. |
|  | Total Number of Spikes |  | NA | The total number of spikes per neuron. |
|  | Total Axon Length |  | μm | Mean total axon length for a neuron. |
|  | Longest branch length |  | μm | Mean for the longest branch length in a neuron |
|  | Longest distance from initiation site |  | μm | Mean for the longest distance from an axon potential's initiation site in a neuron. |
|  | Longest latency from initiation site |  | ms | The amount of time for an action potential to propagate from the initiation site to the beginning of the neuron. |
|  | Conduction Velocity |  | m/s | The velocity of action potential propagation along the neuron. Velocity is the weighted average of the branch velocities based on branch length. |
|  | Amplitude at Initiation Site |  | μV | The peak spike amplitude at the initiation site. |

### Concentration-response modeling with tcplfit2

HdMEA data was then analyzed for each of the 32 endpoints. First, an area under the curve analysis (AUC) was computed for each endpoint to reduce the data by integrating across all DOPs via R statistical computing language (Figure 3). For the AUC analysis method, DOP data were condensed via trapezoidal integration using the *pracma* package (version 2.4.4) in R. Chemically-treated BrainSphere activity was then normalized to DMSO controls by plate-type (6 versus 24-well hdMEA plates), and by endpoint. For normalization, vehicle control wells were pooled, and the median was taken yielding the baseline value (bval) for each plate-type, endpoint, and DOP (variation by plate type is shown in Supplemental Figure 1). Response values from chemically-treated wells were normalized to bval from the same plate type to calculate normalized responses by plate type. The baseline median absolute deviation (bmad) of vehicle controls was computed for each endpoint and an activity threshold cutoff was set to two times the bmad per endpoint.

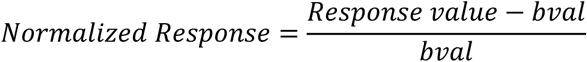

**Figure 3.**
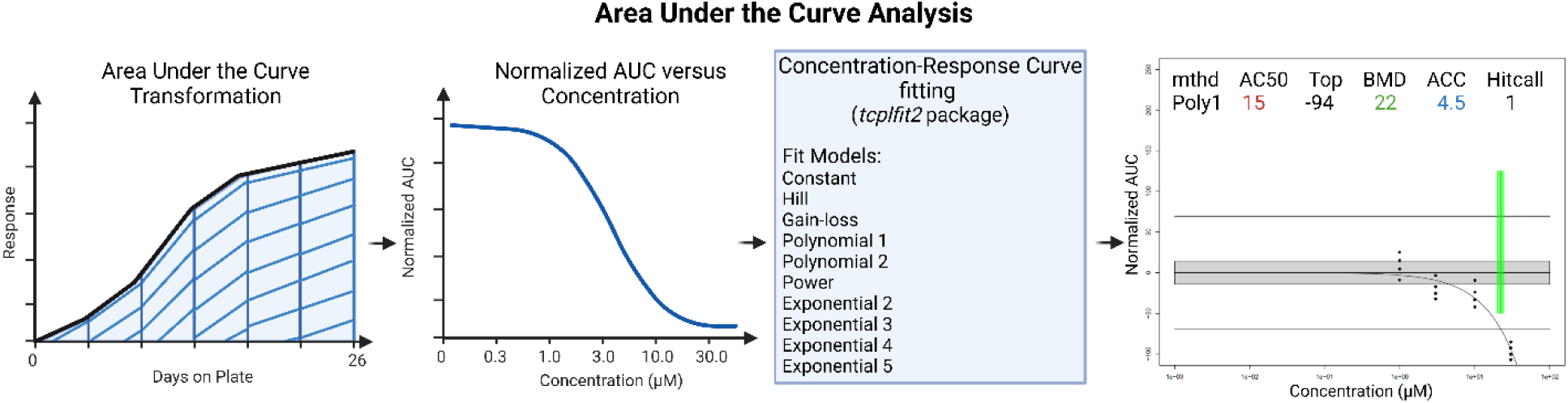
Data Analysis Pipeline. The Area under the Curve (AUC) for response versus days on plate was calculated via trapezoidal transformation using the *pracma* R package resulting in a normalized AUC versus concentration. The normalized results were fit to one of ten models using the *tcplfit2* R package. Chemical-endpoint curves were considered “active hits” if the Hitcall was greater than 0.9. The directionality of the hit was determined by the sign of “Top” using *tcplfit2*.

Concentration-response modeling was performed using the *tcplfit2* package (version 3.1.0) (10.32614/CRAN.package.tcplfit2). Active curves were defined as endpoint-chemical curves that have a continuous hitcall (0 to 1) of ≥ 0.9. Potency was determined using AC_50_ metric, which is defined as the concentration at 50% maximal activity. The log_10_ of the AC_50_ values for each endpoint and chemical were calculated for active curves (Figure 3). The directionality of active chemical-endpoint curves or hits was determined by the sign of “Top”, where the “Top” metric identifies the value at the top of the winning-model curve. All code and source files can be found using the GitHub link: https://github.com/USEPA/Boyd_et_al_2025.

For cytotoxicity data (CellTiter-Blue® and LDH assays), raw values from the plate neutral assay controls were subtracted from each plate from the raw values of the experimental data to eliminate background noise in the assay. Next data was normalized as percent of control using the same equation listed above for hdMEA normalization. Following normalization, CellTiter-Blue® and LDH data was concentration-response modeled using *tcplfit2.* Cutoff values were set to the historical cutoff values from CellTiter-Blue® and LDH assays in the NFA (historical values were unavailable for hdMEA data because the NFA is based on a different MEA system, and therefore the endpoints are not directly analogous). The cutoff values for CellTiter-Blue® and LDH were set to 19.2 and 24.2, respectively. Active curves were defined as endpoint-chemical curves that have a continuous hitcall (0 to 1) of ≥ 0.9.

### Washout Data Analysis

Chemical washout effects on activity were investigated by washing each well twice with fresh medium without chemical treatment and by finally replacing the media with fresh medium without chemical treatment on DOP27. On DOP28 and 29, data were collected (general activity, neural network, and action potential propagation). The activity from the last recording prior to the washout (General Activity and neural network recordings on DOP26 and Action Potential Propagation on DOP27) were statistically compared to the activity following the chemical washout. Raw values were normalized to controls by plate type (24 versus 6-well hdMEA plate), endpoint, and by their respective recording date. The highest two concentrations with measurable endpoints on the day prior to the washout were included for the Wilcoxon test (Dieldrin and deltamethrin= 30 and 100µM, Glyphosate = 10 and 30µM, and loperamide=0.3 and 0.1µM). A paired Wilcoxon test was computed to compare the spontaneous electrical activity prior to washout versus after the washout for every chemical and endpoint with a p-value set to 0.05.

## Results

### Developmental ontogeny of neural activity from the hdMEA 3D BrainSphere assay (untreated)

BrainSpheres were initially characterized to visualize general phenotypic trends in expression of immunocytochemical markers of development at three (timepoint for BrainSphere plating on the hdMEAs) and seven weeks (timepoint for the end of the BrainSphere assay) PID. BrainSpheres at 3-weeks PID express PAX6 and β3-tubulin immunoreactive cells (Supplemental Figure 3). In contrast, BrainSpheres at 7 weeks have PAX6 immunoreactivity, but to a lower extent than week three BrainSpheres. These results are used to qualitatively compare PAX and β3-tubulin immunoreactivity to previous publications to ensure consistency within the model, rather than quantification of protein levels.

Analysis was also conducted to determine the plating time of (3-10 weeks PID) that would result in robust ontogeny of network development and be most representative of the critical period of neurodevelopment. BrainSpheres were plated on hdMEAs at several timepoints ranging from 3-10 weeks PID and grown for approximately 33 days. Qualitatively, the time period of three weeks PID showed robust developmental increases in electrical activity over the course of 33 days, indicative of a phase resembling neural network formation in BrainSpheres (Supplemental Figure 4).

The ontogeny of untreated BrainSphere activity was qualitatively characterized over the course of 28 days to show general trends in BrainSpheres plated at 3 weeks PID while growing on hdMEAs. General activity analysis in control spheroids indicated an initial developmental increase in active area, firing rate (Hz), spike amplitude (µV), and a decrease in inter-spike-interval (ms) over time (Figure 4A; Supplemental Figure 5). These general activity endpoints plateau near the end of ontogeny analysis, suggesting that network activity in BrainSphere may stabilize, potentially indicating a functionally “mature” state. Similarly, endpoints relating to neural network formation (number of spikes per burst and burst peak (Hz)) also resulted in a developmental increase over the course of 28 days growing on the hdMEA (Figure 4B, Supplemental Figure 6). The ontogeny of endpoints relating to action potential propagation were measured and showed a persistent increase in each endpoint across 29 days including firing rate per neuron (Hz) and the number of spikes per neuron (Figure 4C, Supplemental Figure 7).

**Figure 4.**
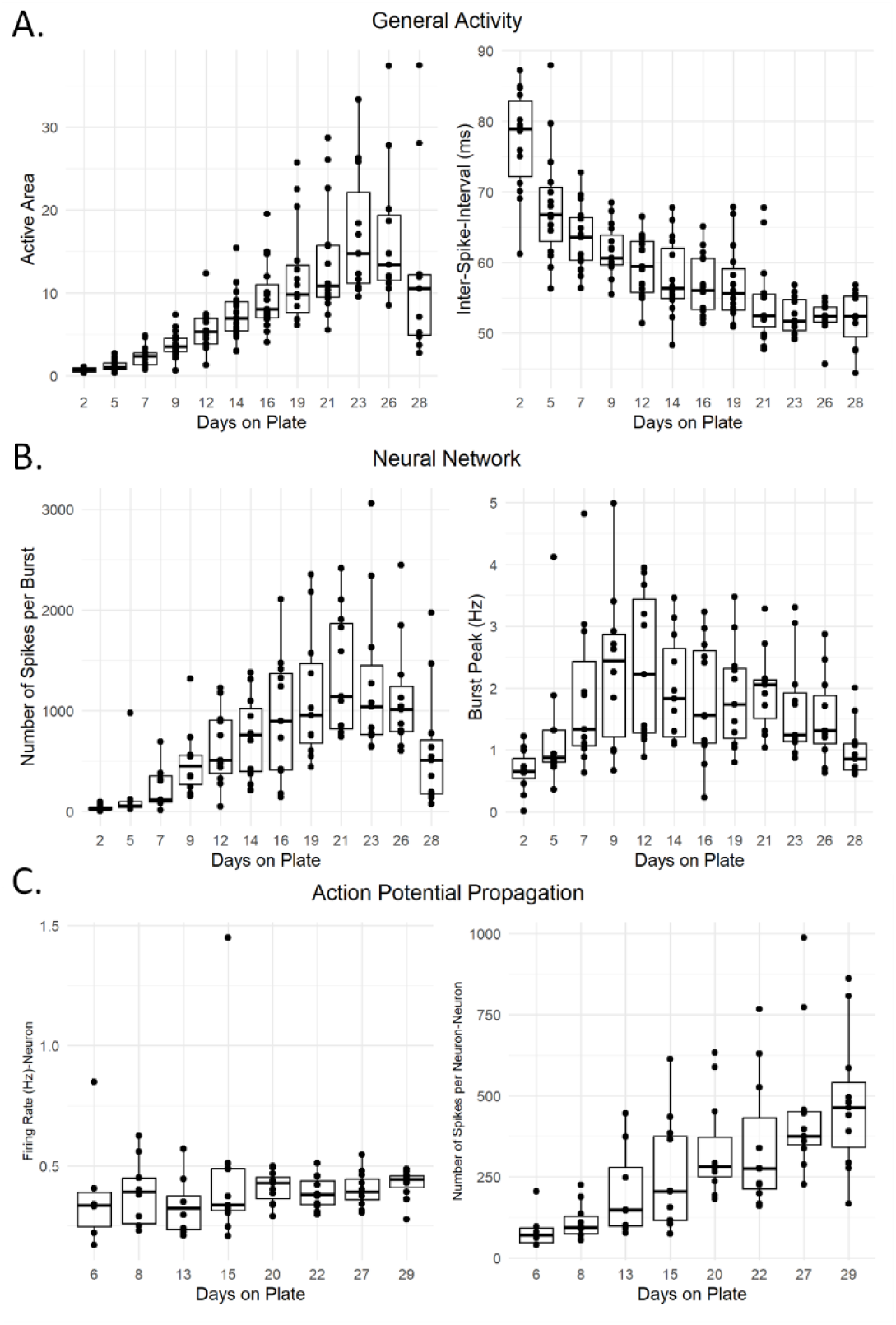
Untreated BrainSpheres growing on hdMEAs demonstrate developmental increases in metrics relating to general activity, neural network, and action potential propagation. A) The ontogeny of general activity for untreated BrainSpheres. The left figure shows active area, and the right represents inter-spike-interval (ms) over the course of 28 days. B) The ontogeny of neural network activity for untreated BrainSpheres. The left figure shows number of spikes per burst, and the right represents burst peak (Hz) over the course of 28 days. C) The ontogeny of action potential propagation endpoints. The mean firing rate represented has Hz per neuron in untreated BrainSpheres on the left and the right image show the number of spikes per neuron on a neuron level basis over the course of 29 days. The data represents an n of 11 BrainSpheres over a total of five hdMEA plates.

When comparing the trends in ontogeny between the different activity types, endpoints relating to neural network activity appear to develop the quickest (peaking near DOP16), whereas general activity peaks near DOP 23, and endpoints of action potential propagation peak closer to DOP29. In addition to characterizing the ontogeny of BrainSphere activity, immunocytochemistry was also conducted to reveal phenotypic characterization throughout BrainSphere development (Supplemental Figure 1). Results show comparable expression levels to previous characterization of the BrainSphere model, indicating a developmental decrease in stem cell markers as well as a developmental increase in mature neural markers (Pamies et al., 2017). Based on the data here and from previous publications, BrainSpheres were plated on hdMEAs three weeks PID for all remaining experiments.

### Technical characterization of the hdMEA BrainSphere endpoints

To perform technical characterization for each endpoint, the CV of control values per endpoint was calculated using the AUC values for each replicate. On average, the CVs for the 32 endpoints in the assay were 27.1% with a range of 4.4-54.8% (Table 2; Supplemental Figure 8). When considering the individual activity types, general activity endpoints had an average CV of 13.1% (6.3-24.6%), neural network endpoints resulted in an average CV of 36.0% (25.1-48.7%), and axon potential propagation endpoints had an average of 25.8% (4.4-54.8%). Evaluating the distribution of CVs across all endpoints (Supplemental Figure 8), we identified 25 endpoints with a relatively lower CV (less than 40%), suggesting that these endpoints may provide greater precision and consistency for measuring activity compared to the 7 endpoints with CVs greater than 40%.

**Table 2.** Coefficient of Variation for each Endpoint in the BrainSphere/hdMEA Assay.

| Recording Type | Endpoint | Level of Data | Coefficient of Variation (%)** |
| --- | --- | --- | --- |
| **General Activity** | Active Area | Well | 24.6 |
|  | Inter-Spike-Interval |  | 6.3 |
|  | Spike Amplitude |  | 9.3 |
|  | Firing Rate |  | 12.1 |
| **Neural Network** | Number of Spikes per Burst | Well | 46.9* |
|  | Inter-Burst-Interval |  | 38.1 |
|  | Burst Peak |  | 42.0* |
|  | Burst Frequency |  | 33.0 |
|  | Burst Duration |  | 25.2 |
|  | Number of Bursts per Cluster |  | 25.1 |
|  | Cluster Frequency |  | 48.7* |
|  | Inter-Cluster-Interval |  | 39.5 |
|  | Cluster Duration |  | 25.2 |
| **Action Potential Propagation** | Branch Length | Branch | 43.5* |
|  | Conduction Velocity |  | 21.5 |
|  | Number of Spikes per configuration |  | 30.9 |
|  | Firing Rate |  | 31.6 |
|  | Conduction velocity coefficient of determination |  | 5.8 |
|  | Longest latency from initiation site |  | 14.9 |
|  | Longest distance from initiation site |  | 44.1* |
|  | Fisher projection distance |  | 13.9 |
|  | Silhouette Score |  | 16.5 |
|  | Total Number of Spikes |  | 30.2 |
|  | Completeness Score |  | 31.5 |
|  | Firing rate | Neuron | 12.1 |
|  | Total number of spikes |  | 24.2 |
|  | Longest branch length |  | 43.5* |
|  | Longest distance from initiation site |  | 23.3 |
|  | Longest latency |  | 31.0 |
|  | Total Axon length |  | 4.4 |
|  | Conduction Velocity |  | 12.4 |
|  | Amplitude at Initiation Site |  | 54.8* |

\*Indicates endpoints that have a CV above 40.0%
\*\* The data represents an n of 11 BrainSpheres over a total of five hdMEA plates.

### Developmental ontogeny of neural activity from dieldrin, deltamethrin, loperamide, and glyphosate-treated 3D BrainSpheres

To determine whether chemical effects on network formation could be measured, BrainSpheres were exposed to assay positive controls dieldrin, deltamethrin and loperamide beginning 2 days after plating on hdMEAs. On DOP29, CellTiter-Blue® and total LDH assays were conducted after the final recording to determine if these treatments impacted cell viability. Data was concentration-response modeled, and zero active curve-fits were identified meaning that no bioactivity was detected for either CellTiter-Blue® or LDH assay for any chemical and concentration tested (Supplemental Files 1-2).

To evaluate compound effects on network formation, each endpoint for general activity, neural network, and action potential propagation analysis were normalized to the vehicle control compounds per DOP and plotted across the days on plate to show the developmental trajectory of chemically treated BrainSpheres grown on hdMEAs. These data were further analyzed using concentration-response modeling to determine active curve-fits for each endpoint and chemical (Figure 6). However, to qualitatively assess trends in chemical perturbations over time, developmental plots were generated (Figure 5). Qualitatively, several endpoints were disrupted as a result of deltamethrin, dieldrin, and loperamide treatment. A representative plot from each type of activity (i.e. general activity, neural network, and action potential propagation) are shown in line graphs for each of the four chemicals (Figure 5), data was further analyzed for changes in biological activity using concentration-response modeling with tcplfit2 (see Figure 6 for concentration-response hits and Supplemental Files 5-7 for concentration-response curves).

**Figure 5.**
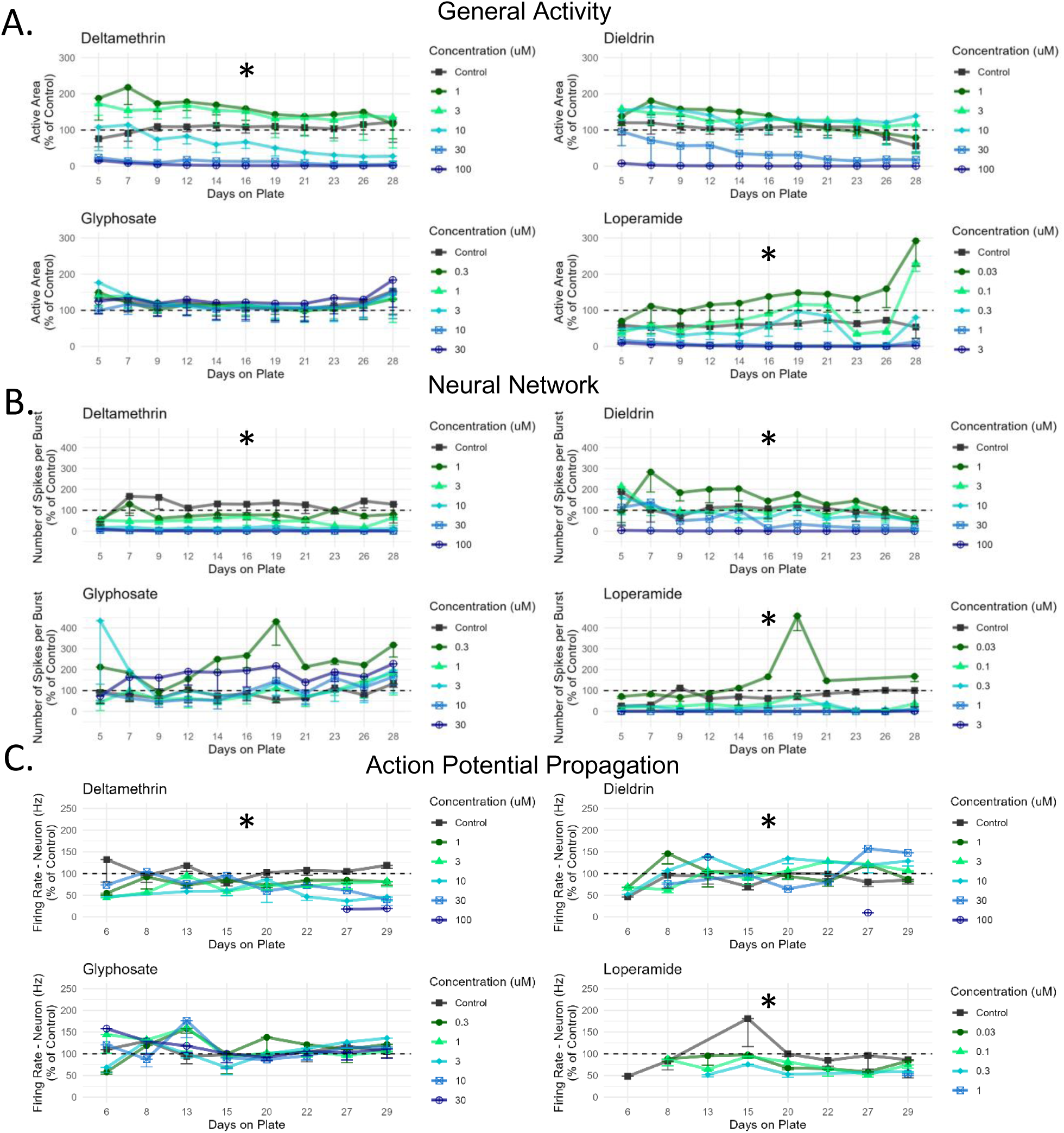
BrainSpheres treated with Deltamethrin, Dieldrin, and Loperamide result in disrupted general activity, neural network, and metrics of action potential propagation. A) Representative line graphs showing active area as percent of control over the course of 28 days on the hdMEA plate. B) Representative line graphs showing number of spikes per neuron a as percent of control over the course of 29 days on the hdMEA plate. C) Representative line graphs showing the firing rate per neuron as percent of control over the course of 29 days on the hdMEA. Error bars represent the standard error and data points show the mean of the replicates. All chemicals and endpoints were concentration-response modeled, and all active curve-fits are indicated with an *. Data represents an n of four for each concentration of dieldrin, deltamethrin, and glyphosate over four hdMEA plates and an n of three BrainSpheres for each concentration of loperamide over three hdMEA plates. See Supplemental Files 3-5 for concentration-response curves and hitcall values.

**Figure 6.**
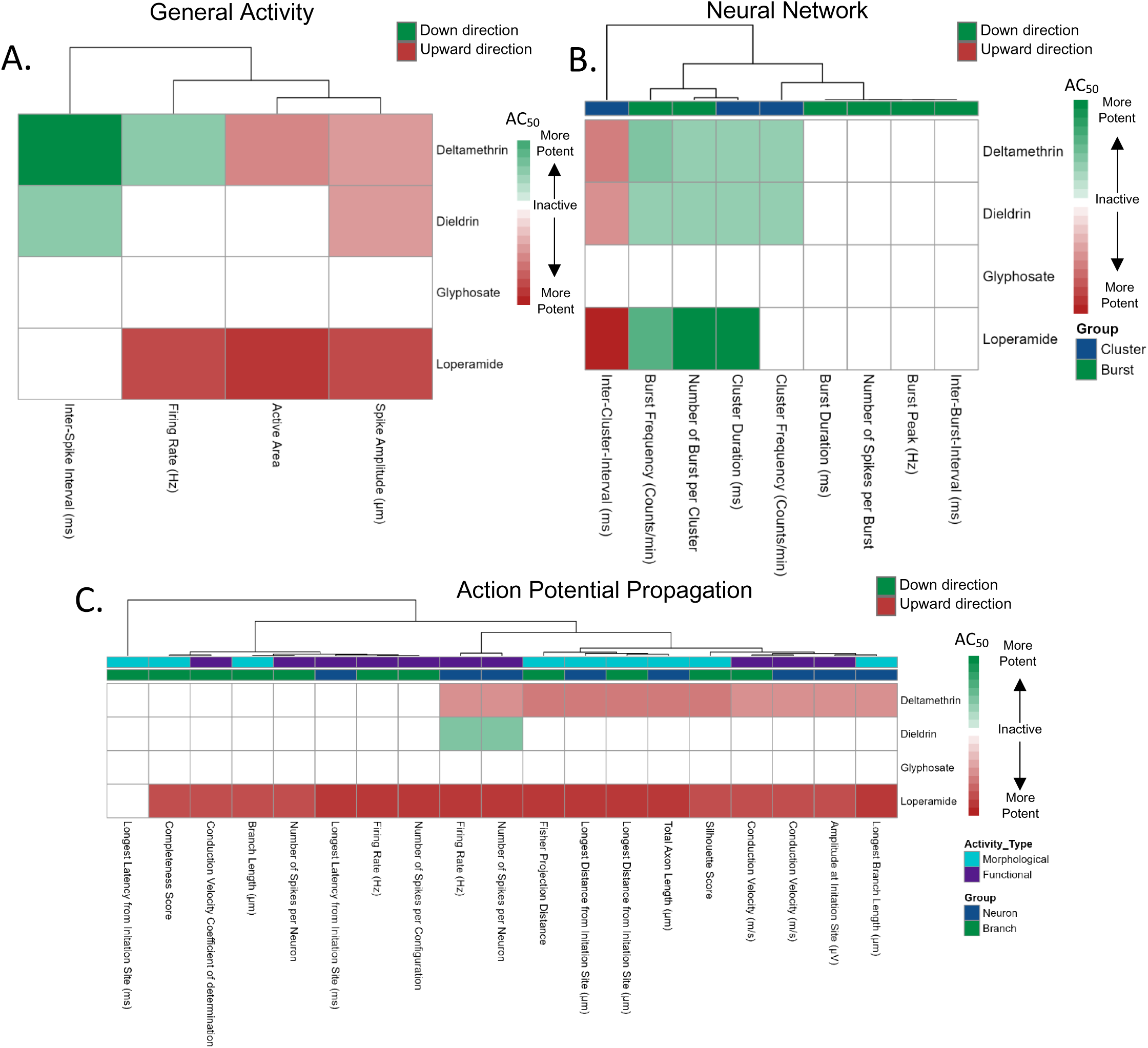
**Area Under the Curve (AUC) analysis indicating th**e **AC50 for all active chemical-endpoint curves for endpoints relating to:** A) General activity B) Neural network-related activity and C) Action potential propagation. BrainSpheres with inactive curves are shown in white and active curve fits are either shown in green or red based on the direction of the curve-fit. Red represents active curve-fits in the down direction while green represents active curve-fits in the up direction. The shade of the color is reflective of the AC50 value, regardless of the direction of the curve-fit. A more potent chemical (smaller AC50 value) would be a darker in color whereas a less potent chemical (larger AC50 value) would be a lighter color. The AC50 values are show in each well of the heatmap and are expressed as µM. Data represents an n of four for each concentration of dieldrin, deltamethrin, and glyphosate over four hdMEA plates and an n of three BrainSpheres for each concentration of loperamide over three hdMEA plates. See Supplemental File 6 for AC50 values.

For general activity, the active area endpoint was shown for each of the four compounds revealing concentration-dependent responses in deltamethrin, dieldrin, and loperamide-treated BrainSpheres (Figure 5A, See Supplemental File 3 for concentration-response modeling results). The low to mid concentrations of dieldrin and deltamethrin-treated BrainSpheres resulted in a steady concentration-dependent decrease in active electrode area over time. In contrast, loperamide resulted in an almost immediate decrease in active electrode area at high concentrations with steady increases in area for low concentrations of loperamide (0.03µM). Regarding neural network activity, the number of spikes per burst is shown as a representative endpoint for each of the four chemicals tested, demonstrating concentration-dependent disruptions for deltamethrin, dieldrin, and loperamide treated BrainSpheres (Figure 5B; See Supplemental File 4 for concentration-response modeling results). Mid to high concentrations of deltamethrin and loperamide resulted in an almost immediate, concentration-dependent decrease in the number of spikes per burst. Mid to low concentrations of dieldrin resulted in a steadier, concentration-dependent decrease in the number of spikes per burst. The lowest concentrations of loperamide and dieldrin resulted in increases in the number of spikes per burst above baseline, and the peak of this effect occurred later in loperamide-treated BrainSpheres. Firing rate on the neuron level is used as a representation for action potential propagation (Figure 5C; See Supplemental File 5 for concentration-response modeling results). Dieldrin, deltamethrin and loperamide-treatments resulted in concentration-dependent disruption in firing rate, although these disruptions are less apparent than some of the general activity and neural network-based endpoints.

Deltamethrin and loperamide both resulted in a steady concentration-dependent decrease in firing rate over DOPs in the absence of cytotoxicity. In contrast, dieldrin-treated BrainSpheres resulted in a steady increase in firing rate over DOPs, most notably with high concentrations. Each endpoint and chemical was then concentration-response modeled and active curve-fits are described below (Figure 6).

### Concentration-response modeling detected chemical-induced effects

Dieldrin, deltamethrin, and loperamide treatments resulted in a total of nine curve-fits with a positive hit call determination (‘bioactive’) for endpoints relating to general activity (Figure 6A, See Supplemental File 6 for corresponding AC50 and hitcall values). Further analysis reveals that three out of the nine bioactive curves for general activity were considered active in the up direction for inter-spike interval, indicating an increase in the inter-spike interval and thus suggesting a decrease in neural spiking activity. However, one hit for firing rate was detected in deltamethrin-treated BrainSpheres indicative of increased firing activity. The remaining curve-fits detected activity in the down direction for firing rate, spike amplitude, and active area, suggesting decreasing neural activity in the assay for the three chemicals. Glyphosate, the assay negative control, did not result in any hits for endpoints relating to general activity (Figure 6A), indicating a lack of effects on bioactivity.

Dieldrin, deltamethrin, and loperamide disrupted several endpoints relating to neural network activity in the increased and decreased direction (Figure 6B, See Supplemental File 6 for corresponding AC50 and hitcall values). Deltamethrin and dieldrin resulted in the greatest number of hits across all neural network endpoints with a total of five bioactive curves out of the nine possible endpoints, followed by loperamide with four hits (Figure 6B). The curve-fits detected bioactivity in the up direction for cluster frequency, cluster duration, burst frequency, and the number of bursts per cluster. The upward direction of these endpoints indicates an increase in the AUC, or an increase in bioactivity relative to the DOPs. For example, an increase in the AUC of the cluster frequency endpoint, may indicate an increase in cluster formation across time resulting in a greater AUC value. The remaining curve-fits detected activity in the down direction for the remainder of the neural network endpoints. Glyphosate did not induce any curve-fits with bioactivity (Figure 6B).

For endpoints relating to action potential propagation, several curve-fits were active for dieldrin, deltamethrin, and loperamide, while glyphosate exposure was inactive (Figure 6C, See Supplemental File 6 for corresponding AC50 and hitcall values). Out of 19 endpoints, loperamide resulted in the greatest number of hits (18), followed by deltamethrin (11) and then dieldrin (2). Two endpoints (firing rate-neuron and the number of spikes per neuron) resulted in an increase in bioactivity rather than a decrease, all of which occurred in dieldrin-treated BrainSpheres (Figure 6C).

### Chemical Washout

After the last hdMEA recording on DOP27, a washout was performed for each well by exchanging the medium twice with fresh medium as a wash and finally replaced with fresh, chemical-free differentiation medium for the remainder of the experiment. This washout was used to determine whether effects may be explained by acute exposure in the assay (last chemical addition on DOP 26). Activity on the day prior to and after the washout (DOP 26 versus DOP 28) were compared for each endpoint, treatment, and concentration (Supplemental File 7). Only the effects of deltamethrin for the general activity endpoint ISI was significantly affected (p-value of 0.003), suggesting that ISI may have been perturbed following the washout. Prior to the washout, deltamethrin resulted in an active curve-fit in the upward direction for ISI, meaning that ISI was increased with deltamethrin treatment. However, with the washout deltamethrin’s ISI was decreased. ISI has a low coefficient of variation (CV) of 6.3% indicating that the washout effect observed here for deltamethrin is not due to high variability in this endpoint (Table 2). However, in total only one significant value was identified out of a total of 128, indicating that a chemical washout did not result in any widespread recovery of spontaneous electrical activity in chemically treated BrainSpheres. This indicates that effects of the chemicals are sustained and long-lasting and likely do not represent acute pharmacological effects.

## Discussion

Demonstrated here is a complex 3D BrainSphere model which offers promise for investigating toxicological perturbations to the development of neural networks. First, the ontogeny of electrical activity was characterized indicating a developmental increase in electrical activity in BrainSpheres as they are growing on hdMEAs for 29 days. Thirty-two unique endpoints were used to describe BrainSphere general activity, neural network formation, and action potential propagation. For the current study, dieldrin, deltamethrin, and loperamide were selected as positive control compounds since these chemicals induce known disruptions in neural network formation in primary rat cortical neurons (Carstens et al., 2022). Glyphosate was selected as an “intermediate” negative control chemical because of the lack of DNT-related effects demonstrated in the primary rat cortical network formation assay (Martin et al., 2022; Carstens et al., 2022). Glyphosate was considered an “intermediate” negative control rather than a “favorable” negative control due to bioavailability concerns, however, glyphosate proves to be inactive in the rat NFA, making it an ideal negative compound for the current assay (Martin et al., 2022; Carstens et al., 2022). This subset of training chemicals was tested in the BrainSphere assay resulting in expected toxicological perturbations to network development paralleling previous results from the rat NFA for the same chemical set (Carstens et al., 2022). The assay also yielded additional data beyond what is offered by the NFA such as the action potential propagation endpoints. Such information may be useful for understanding a chemical’s mechanism of action. These toxicological effects occurred in the absence of cytotoxicity and the effects were sustained following a chemical washout. Therefore, the assay described here proves to be a functional, complex, sensitive, and human relevant model that can be used to detect alterations in network formation induced by chemical exposure.

When developing an assay aimed at modeling a key event of neurodevelopment, like neural network formation, it is essential to establish a model that accurately and robustly recapitulates the features and characteristics essential for that key neurodevelopmental event (Crofton et al., 2011; OECD, 2023).

Previously, the primary rat cortical MEA-based assay (NFA) has undergone extensive characterization of neural activity captured by the assay (Brown et al., 2016). Prior work has shown that the NFA is sensitive to toxicological perturbations induced by a training set of DNT chemicals and that the 17 endpoints describe network connectivity, burst structure, and synchrony. The experimental paradigm of the NFA has also been characterized, demonstrating that the 2D rat cortical model captures increase in activity over the course of two weeks indicating recapitulation of neural network development and formation (Brown et al., 2016). In the current project, extensive characterization of the ontogeny of the activity of BrainSpheres plated on hdMEAs was conducted at timepoints three to ten weeks post-initiation of differentiation. Analysis revealed that BrainSpheres plated at three weeks PID showed the greatest increases in activity over the course of 29 days grown on the hdMEAs indicative of a robust, developmental neural network model (Supplemental Figure 4). Supplementing this work, BrainSpheres were also characterized for general trends in the expression of PAX6 (a marker of neural progenitor cells) and β3-tubulin (a marker of mature neurons) (Supplemental Figure 3). Results indicated that PAX6 expression decreased as the BrainSpheres developed between weeks three and seven post-initiation of differentiation, demonstrating that the selected timepoints captured developmental decreases in neural progenitor cell immunoreactivity. This result is consistent with previous work characterized phenotypic ontogeny of BrainSpheres showing that BrainSpheres have RNA expression for markers of GABAergic, glutamatergic, and dopaminergic neurons (Pamies et al., 2017).

To characterize the ontogeny of spontaneous electrical activity in the time period captured by the current assay, three-week BrainSpheres were plated on hdMEAs to recapitulate a developmental phase of neural networks. Endpoints relating to general activity and neural network exhibited a time-dependent increase in activity followed by a plateau in activity (Figure 4A-B; Supplemental Figures 5 and 6). However, the ontogeny of action potential propagation-related endpoints appears to reach its peak, in terms of activity, several days later than general activity and neural network-related endpoints (Figure 4C; Supplemental Figure 7). The differences in the time to peak levels of activity between the types of activity recorded (i.e. general activity, neural network, and action potential propagation) highlights that each of these three features of network development and neuronal communication may occur on different time courses with unique trajectories. Toxicological perturbations may disrupt the developmental trajectory, impacting the time to peak effect for each of these features during network formation. These activity trajectories may track with phenotypic characterization observed previously in the model (Pamies et al., 2017).

When investigating the variability of each endpoint, CVs were computed for each individual endpoint using robust statistical measures (median absolute deviation/ median of controls). Previous reports observed that endpoints measuring neural activity in an MEA-based assay had an average CV of 18.23

+/-6.15% (2D rat cortical neurons on standard MEAs) (Carstens et al., 2022) or 16.96 +/-8.21% (Human iPSC-dervied SynFire model on standard MEAS) (Bartmann et al., 2023) or 62.4 +/-27.1% (hdMEA BrainSphere model using a human iPSC cell model) (Carstens et al., 2025). Notably, in Carstens et al., 2025, the assay evaluated a complex 3D BrainSphere model representing a more mature phase biologically (7-week BrainSpheres at plating), where multiple BrainSpheres were grown in one well of 6-well plates with a limited number of vehicle controls, which may explain high average CVs.

The current assay resulted in 25/32 endpoints that demonstrated a CV below 40%, suggesting these endpoints may be more reliable and reproducible for chemical screening compared to the remaining 7 endpoints with higher CVs. The CVs for endpoints relating to neural network activity were the highest with 3/9 endpoints greater than 40%. Based on the results of this study it appears that the development of neural network activity may be more variable compared to general activity. A total of 4/19 action potential propagation endpoints had CVs greater than 40%, and 3/4 of the endpoints were morphologically based. The higher CV for the morphological action potential propagation endpoints such as branch length compared to the functional action potential propagation endpoints may be partly related to the 3D structure of the BrainSphere which may cause an inability to detect the full axonal processes in BrainSpheres since axons may grow in the Z-plane, which the MEAs cannot detect.

Additionally, many endpoints relating to action potential propagation are undetectable at early recording days, and as the BrainSpheres mature additional endpoints become detectable at later recording days. Although the variability in baseline control values were slightly higher in the current assay than some of the comparable monolayer MEA-based assays, the variability was expected to be higher considering the complexity of the BrainSphere structure, the length of the assay, and the higher temporal and spatial resolution of the hdMEA system compared to the standard MEA system.

For the high-density MEA assay, three types of recording modules were conducted to capture general activity, neural network activity, and action potential propagation (Figure 2). Many of the endpoints provided by the high-density system are not directly comparable to those provided by the standard density MEA systems. This is especially true for the action potential propagation endpoints which provide additional value to the assay by describing both morphological and functional features of action potential propagation. Disruption to these endpoints might provide mechanistic insight on chemical perturbation to network formation. For example, dieldrin resulted in a bioactive curve-fit for firing rate on the neuron level (Figure 6C). The acute mechanism of action for dieldrin is well established and is known to inhibit the chloride influx through GABAA receptors, therefore resulting in suppression of inhibitory neurons. In the current assay, dieldrin results in a bioactive curve-fit in the upward direction for firing rate (Figure 6). This directional effect indicates that the area under the curve for firing rate over the time course is greater than vehicle treated BrainSpheres showing an overall increase in neuronal activity. This unique increase in firing rate for dieldrin is captured in the action potential propagation endpoints, providing potential insight on mechanisms for dieldrin disruption of network formation.

These results recapitulate dieldrin-induced bioactive curve-fits in the up direction in the primary rat cortical NFA. Specifically, dieldrin induces increases in the number of spikes in a network spike and spike duration in the NFA (Carstens et al., 2022) (Table 2). Additionally, some of these chemical-specific hits in the action potential propagation endpoints provide added value that would not be captured when only looking at general and neural network-related endpoints, such as in the NFA.

Additionally, morphological endpoints such as the action potential propagation endpoint, longest branch length or the general activity endpoint, active area may be reflective of neurite outgrowth related changes as well. Two assays included in the DNT-IVB investigate neurite outgrowth in human neural progenitor cells (iCell GlutaNeurons™) and primary rat cortical cells (Harrill et al., 2018). Dieldrin and loperamide both affected many of the endpoints included in these neurite outgrowth assays, with loperamide producing effects with a greater potency than dieldrin (Carstens et al., 2022), consistent with the findings of this study (Table 3). Another assay included in the battery is the Synaptogenesis and Maturation Assay which uses primary rat cortical cells to assess synaptic features of complex neural connections (Harrill et al., 2011). Since neurite outgrowth is required for the formation of synapses, synaptogenesis and maturation may also serve as a proxy marker for morphological features of neurodevelopment. Deltamethrin has been shown to affect several synaptogenesis endpoints (Carstens et al., 2022). In relevance to the current assay, dieldrin, deltamethrin, and loperamide all induce toxicological perturbations on active area, which is an endpoint indicative of neurite outgrowth across the electrode surface area over time, with loperamide showing the greatest potency (Figures 6A, 1A).

**Table 3.** Comparison of AC50 values per activity type between 2D and 3D models systems and the current assay.

| NFA<br>2D rat primary cortical cultures/MEA<br>(Carstens et al., 2022) |  |  |  | BrainSphere/hdMEA<br>7 Weeks PID<br>(Carstens et al., 2025) |  |  |  | BrainSphere/hdMEA<br>3 Weeks PID<br>(Current Study) |  |  |  |
| --- | --- | --- | --- | --- | --- | --- | --- | --- | --- | --- | --- |
| Activity Type | Chemical | Mean AC<sub>50</sub> (μM) | Min AC<sub>50</sub> (μM) | Activity Type | Chemical | Mean AC<sub>50</sub> (μM) | Min AC<sub>50</sub> (μM) | Activity Type | Chemical | Mean AC<sub>50</sub> (μM) | Min AC<sub>50</sub> (μM) |
| Neural Network | Dieldrin | 7.75 | 0.70 | General Activity | Dieldrin | 14.33 | 9.60 | General Activity | Dieldrin | 34.455 | 18.91 |
|  | Deltamethrin | 4.62 | 0.61 |  | Deltamethrin | 4.44 | 1.40 |  | Deltamethrin | 28.70 | 0.07 |
|  | Loperamide | 0.93 | 0.21 |  | Loperamide | 0.23 | 0.03 |  | Loperamide | 0.24 | 0.12 |
|  |  |  |  | Neural Network | Dieldrin | 14.14 | 9.45 | Neural Network | Dieldrin | 26.19 | 9.23 |
|  |  |  |  |  | Deltamethrin | 6.4 | 3.49 |  | Deltamethrin | 15.80 | 0.99 |
|  |  |  |  |  | Loperamide | 2.51 | 0.02 |  | Loperamide | 0.12 | 0.004 |
|  |  |  |  |  |  |  |  | Action Potential Propagation | Dieldrin | 70.23 | 50.00 |
|  |  |  |  |  |  |  |  |  | Deltamethrin | 9.42 | 2.35 |
|  |  |  |  |  |  |  |  |  | Loperamide | 0.33 | 0.20 |

Additionally, deltamethrin and loperamide perturb several morphological features of action potential propagation as well such as total axon length or the longest distance from initiation site for example (Figure 6C). Loperamide induces potent and bioactive responses for all eight morphological action potential propagation endpoints. Therefore, the hdMEA-based assay also provides informative measurements that may serve as an orthogonal assay for neurite outgrowth features.

In the current study, loperamide treated-BrainSpheres resulted in 25, deltamethrin (20), dieldrin (9), and glyphosate (zero) hits out of 32 possible endpoints (Figure 6). These results are concordant with comparable neuronal MEA-based models, although it is difficult to directly compare chemical perturbations and potency amongst these different models due to discrepancies in MEA systems and general model differences (human iPSC-derived monolayer and 3D cultures versus monolayer rodent cultures versus standard and high density MEAs) (Carstens et al., 2022; Saavedra et al., 2021; Bartmann et al., 2023; Hartmann et al., 2023; Carstens et al., 2025). Out of these existing models, Carstens and colleagues have reported similar toxicological perturbations to a more mature BrainSphere model (plated at seven weeks PID as opposed to three weeks in the current study) on high-density MEAs using a shorter exposure and recording paradigm (13 days as opposed to four weeks) (Carstens et al., 2025).

Carstens and colleagues demonstrated concentration-dependent disruption in neural network and general activity endpoints for loperamide, dieldrin, and deltamethrin-treated BrainSpheres (Table 3). When comparing the minimum AC50 values, the current BrainSphere assay captures toxicological perturbations with greater sensitivity compared to the BrainSphere assay introduced by Carstens and colleagues (Table 3) (Carstens et al., 2025). Similarly, the current BrainSphere assay also demonstrates greater sensitivity when comparing the minimum AC50 values of the NFA, except for the case of dieldrin (Table3)(Carstens et al., 2022).

In comparison to the BrainSphere assay by Carstens and colleagues, the model introduced here is representative of a more developmental exposure to neuronal networks because of the younger age of the BrainSpheres at plating. Moreover, in the current study we have characterized the ontogeny of BrainSpheres over the course of ten weeks and have identified that plating BrainSpheres at 3-weeks PID results in the greatest developmental increases in activity. Meaning that BrainSpheres result in large developmental changes in spontaneous electrical activity when plated on hdMEAs at 3-weeks PID, which is not reflected when plated at other timepoints PID (Supplemental Figure 4). It is well established that high levels of spontaneous electrical activity occur during critical periods of neurodevelopment, underpinning the rapid development of neurons, glia, and synaptic connections. However, as the brain matures the spontaneous electrical activity declines, and this decline is thought to be due to slowed rates of synaptic formation and synaptic pruning (Banies and Landgraf, 2021; Levitt, 2003, Tau et al., 2009). Therefore, the BrainSphere assay presented here represents an earlier phase of neurodevelopment with high neuronal activity, a time period sensitive to toxicological perturbations (Rock and Patisaul, 2019).

A chemical washout was conducted at the end of the experiment, followed by an additional set of recordings 24 hours after the washout. The washout was used to determine if the perturbations in network function observed here are the result of acute, pharmacological effects, or if these effects reflect long-lasting and sustained impacts on neural networks. Doing so, the recording data prior to the washout was compared to the recording data after the washout resulting in a total of 128 comparisons, of which only one demonstrated a significant change in activity following the washout (Supplemental Table 1). The lack of washout-induced effects is indicative of long-lasting developmental changes, since even in the absence of a chemical readily available in the media, the BrainSpheres continued to show disrupted neuronal activity. However, a limitation of the washout is that additional manipulation occurring during the washout step may interfere with the spontaneous electrical activity of the BrainSpheres. It is also possible that a chemical washout may not completely remove all the chemical in a specific well, particularly for lipophilic chemicals such as deltamethrin which may remain within cells (Shafer et al., 2008). The NFA does not include a washout step in the assay, however others have reported similar washout procedures in MEA-based models (Bartmann et al., 2023; Hartmann et al., 2023).

The DNT-IVB incorporates several NAMs, each providing data informative on one of the essential processes comprising neurodevelopment. Together, data from each assay in the battery provides bioactivity information about chemical effects and potencies on these critical processes that can be used towards hazard identification and/or prioritization (OECD, 2023; Juberg et al., 2023, U.S. Environmental Protection Agency, 2020). Prior expert reviews that provided support for use of the DNT-IVB in these contexts also highlighted that the current battery can be improved upon by developing models that more fully recapitulate the complexity and nuances of the human developing brain (Juberg et al., 2023; U.S. Environmental Protection Agency, 2020).

Addressing the concern about model complexity, a recent publication compared the transcriptomic profiles of human iPSC-derived 2D versus 3D models (Lislien et al., 2025). Lislien and colleagues demonstrated that 3D models have improved differentiation resulting in a more heterogeneous cellular composition and most notably higher expression of GABAergic, dopaminergic, glycinergic, serotonergic, and cholinergic populations. Similarly, there is greater expression of pre-and post-synaptic markers in the 3D cultures compared to 2D cultures (Lislien et al., 2025). Synaptic proteins are essential for forming neuronal connections and ultimately facilitating neuronal network formation (Connors et al., 2008; Batool et al., 2019). Additionally, the period over which human iPSCs develop network activity *in vitro* compared to rodent cells is prolonged in comparison to rat cortical neuronal cultures, more consistent with the long neurodevelopmental period exhibited in humans and increases the biological window for DNT testing (Pamies et al., 2018; Pamies et al., 2014; Hopkins et al., 2015). Overall, these results suggest that 3D human iPSC-derived models serve as a more biologically relevant model of both the heterogeneity and the complexity of neuronal connections observed in the human brain, albeit with the potential restriction of lower assay throughput (Lislien et al., 2025). Therefore, we set forth to build a novel assay based on the NFA, the only functional based assay that is amenable for inclusion in the DNT-IVB (Brown et al., 2016). The NFA utilizes 2D primary rat cortical cells and standard electrode arrays to provide information on toxicological perturbations to neural network formation, a cumulative product of all neurodevelopmental processes such as proliferation, synaptogenesis, neurite outgrowth, etc. and is the underlying molecular basis of high-order brain functioning like cognition (Khazipov and Luhmann, 2006; Mennerick and Zorumski, 2001; Buschman and Kastner. 2015; Salinas and Sejnowski, 2001). The BrainSphere model demonstrated here represents a next generation assay that addresses some limitations of the rat NFA and improves upon its human relevancy, complexity, and functionality.

Considering the cost, lengthy timeline, and labor demands of the BrainSphere assay, a tiered approach testing strategy would best facilitate the use of the assay. Tiered testing provides a framework for more rapid screens to identify chemicals with putative hazard followed by more specific testing strategies which improve on the complexity and human/tissue relevancy (Thomas et al., 2019). In terms of a proposed tiered testing strategy for DNT, a high-content phenotypic profiling approach or high throughput transcriptomics could be employed as a first tier for assessing potential DNT hazard (Culbreth et al., 2022). Chemicals with identified hazard in tier one would be further investigated in tier two, which in this case would be the DNT-IVB since the battery is less labor intensive and expensive than the current assay but still provides important information. Following additional standardization and validation experiments, one could consider that tier three would include the BrainSphere hdMEA assay, demonstrated here to provide further characterization and mechanistic information in a more human-relevant and complex model. Additional assays may also be used to supplement the BrainSphere assay proposed here such as a behavioral assay using Zebrafish such as the photomotor response behavioral assay in larval zebrafish (Truong et al., 2014). As a chemical moves through the different tiers, the assays become more complex and biologically relevant to aspects of neurodevelopmental processes, such as neural networks. Therefore, the current BrainSphere assay has potential to serve as a highly complex system that can be used to provide further hazard characterization as well as gain mechanistic information for DNT chemicals following screening with the DNT-IVB. Future work should be aimed at testing additional chemicals in this new model and further refining of the assay. The current assay was sensitive to detecting chemically induced perturbations in neural network function and action potential propagation and may provide a useful complex tissue model to assess DNT potential of chemicals.

## Acknowledgements

The authors would like to thank Alan Tennant for technical expertise with confocal microscopy, and Kathleen Wallace and Theresa Freudenrich for general laboratory support. Additional acknowledgments go to Katie Paul-Friedman, Kimberly Slentz-Kesler, and Alison Harrill for their constructive feedback and edits on the manuscript.

