## Supplemental Figures for "Development of a network formation assay for developmental neurotoxicity hazard screening using 3D human iPSC derived BrainSpheres"

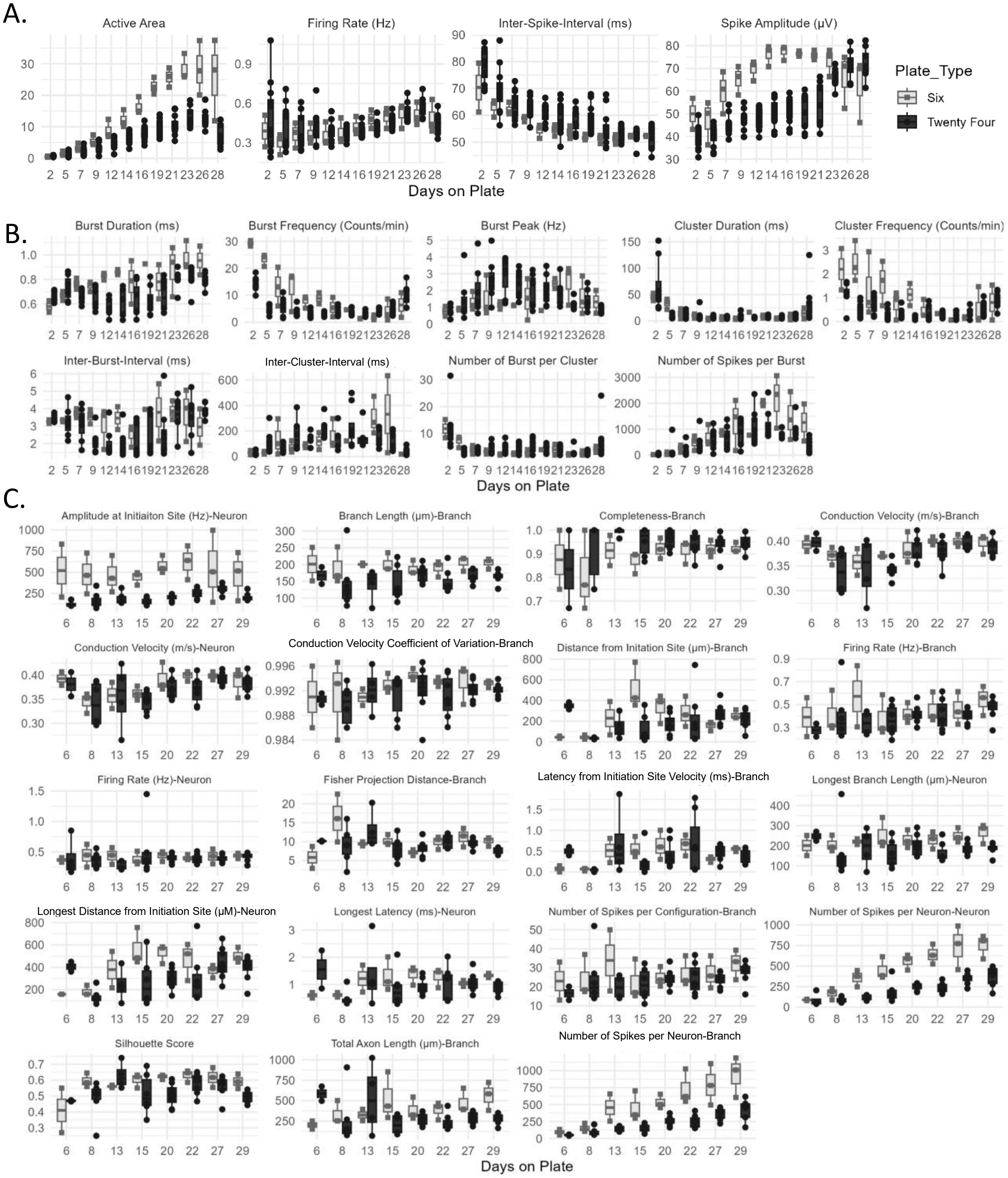


**Supplemental Figure 1. Ontogeny of spontaneous electrical activity in untreated BrainSpheres grown on each type of high-density microelectrode array plate for endpoints of** A) General Activity, B) Neural Network, and C) Action Potential Propagation. Data represents 3, 24-well plates each with 4 BrainSpheres per plate and 3, 6-well plates with 1 BrainSphere per plate.

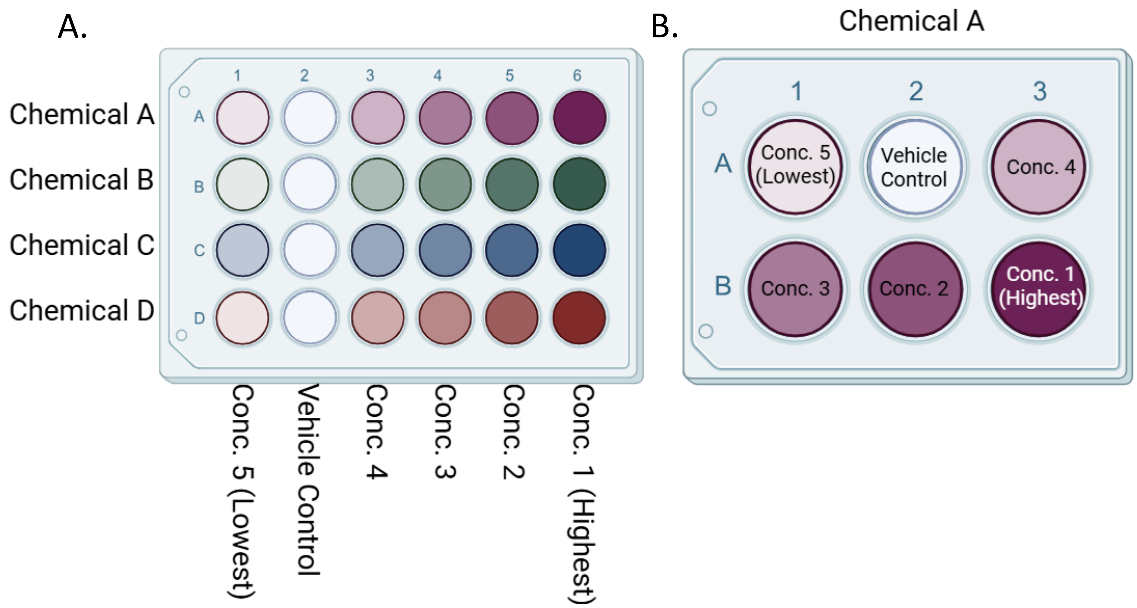


**Supplemental Figure 2. Example hdMEA plate layouts for** A) 24-well and B) 6-well plates. For 24-well hdMEA plates, BrainSpheres were treated with one of four different chemicals, where each row of the plate is treated with a different chemical. Each chemical was tested at five concentrations and a vehicle control. Each row of the plate was treated with a different concentration of the chemical (Conc. 1-5). The highest concentration tested was in column 6, whereas the lowest concentration tested was in column 1. The vehicle controls (DMSO for dieldrin, deltamethrin, and loperamide or water for Glyphosate) were in column 2 to avoid edge effects. For each 24-well plate, the rows (chemicals) were randomized so chemical replicates were tested in a unique row position. For 6-well plates, one chemical was tested per plate at 5 concentrations (Conc. 1-5) and a vehicle control. The highest concentration tested was in well B3 and the lowest concentration tested was in well A1. The vehicle controls were tested in well A2. The remaining doses were tested in wells A3, B1, and B2. Each chemical was tested in a combination of 24-well and 6-well plates due to availability of plates at the time of experimentation.

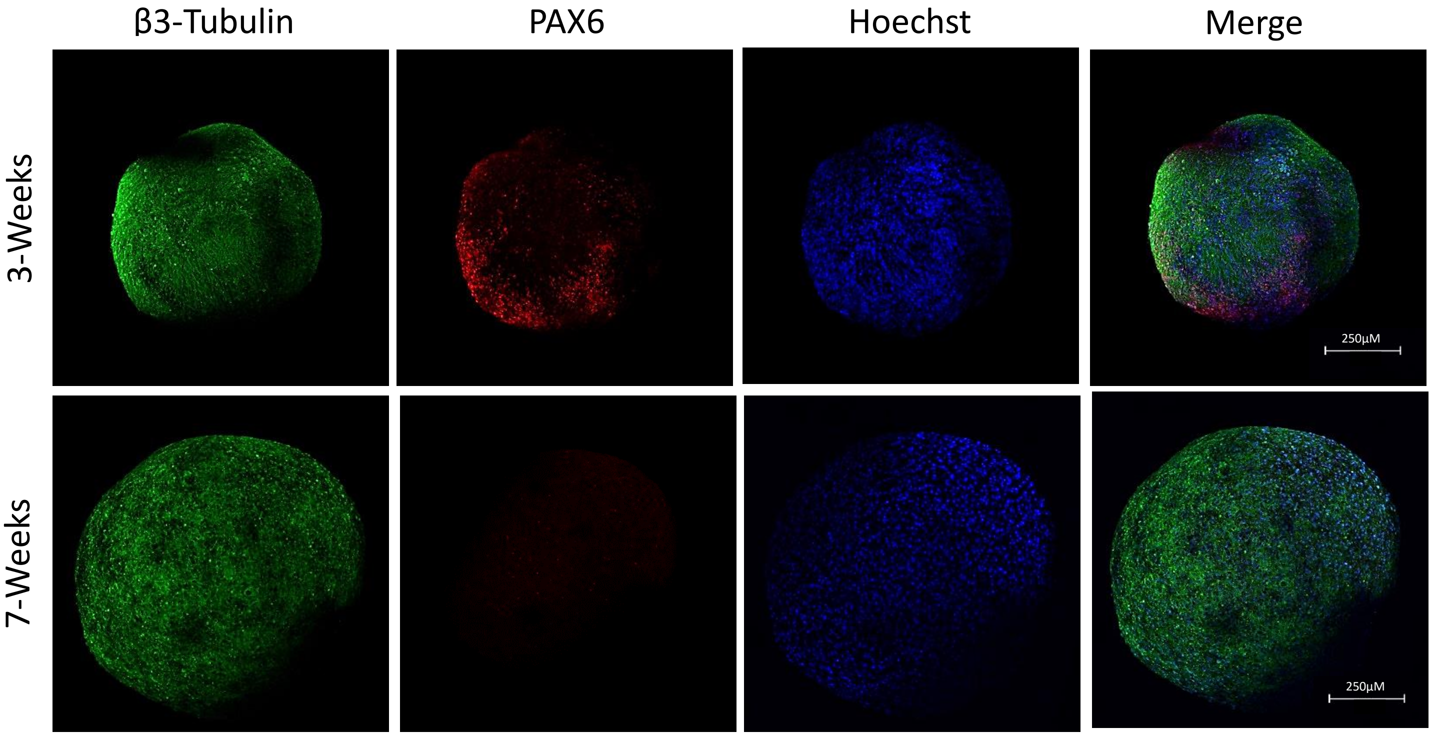


**Supplemental Figure 3. Expression of neural marker, β3-Tubulin and the neural progenitor marker, PAX6 for 3- and 7-weeks post initiation of differentiation in representative BrainSpheres.** Green, red and blue pseudo coloring represents β3-Tubulin,PAX6, and Hoechst immunoreactivity in BrainSpheres 3-weeks post-initiation of differentiation (top panels) and 7-weeks post-initiation of differentiation (bottom panels).

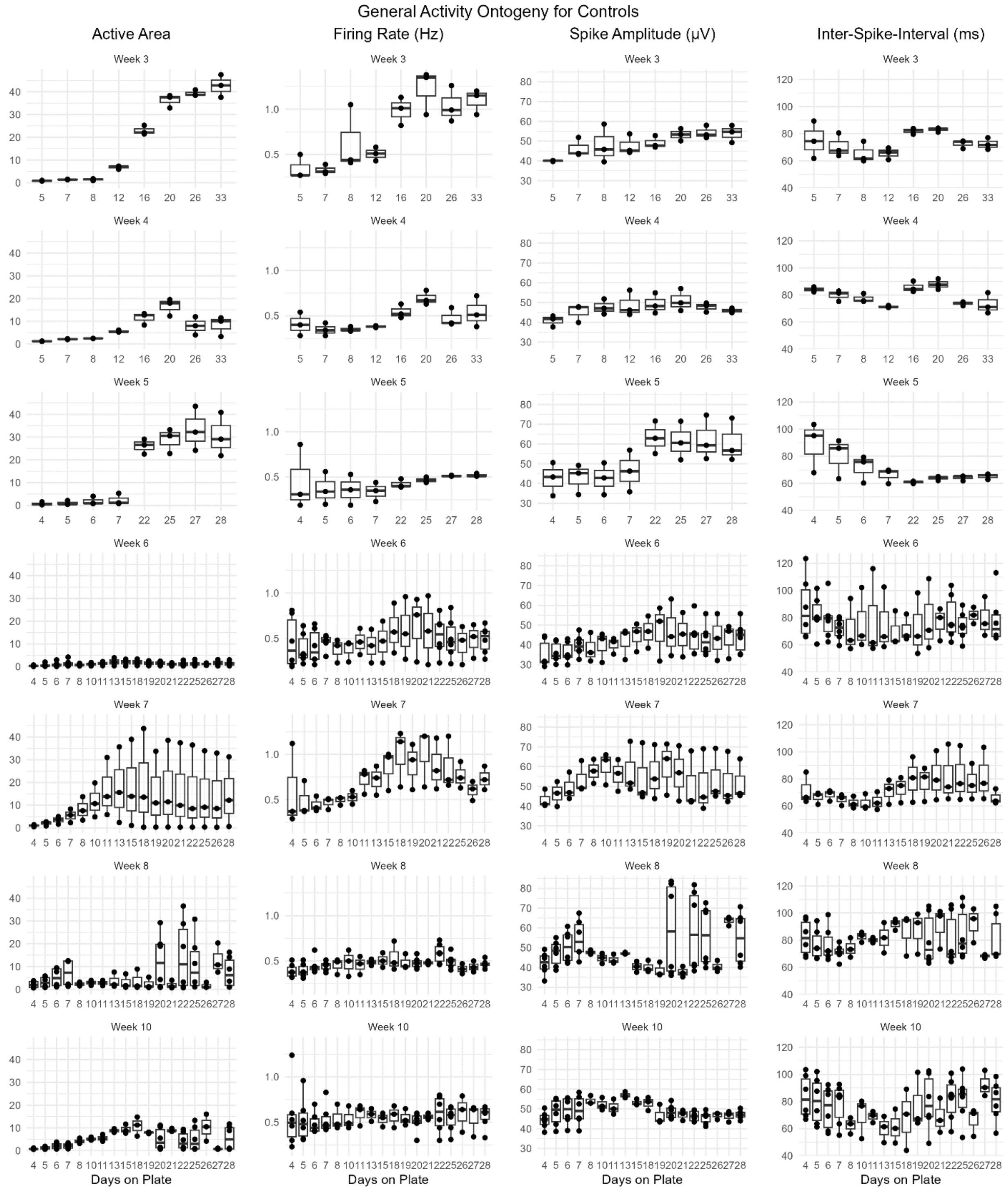


**Supplemental Figure 4. Ontogeny of general activity in control BrainSpheres over 10 weeks.** Graphs represent the number of days on plate across the x-axis for each of the different metrics describing general BrainSphere activity shown on the y-axis for BrainSpheres plated at weeks three, four, five, six, seven, eight, and ten. For plating weeks three, four, and five, one plate was used for each plating week and each plate had an n of three BrainSpheres. For plating weeks six, seven, eight, and ten, two plates were used for each plating week and each plate had an n of three BrainSpheres.

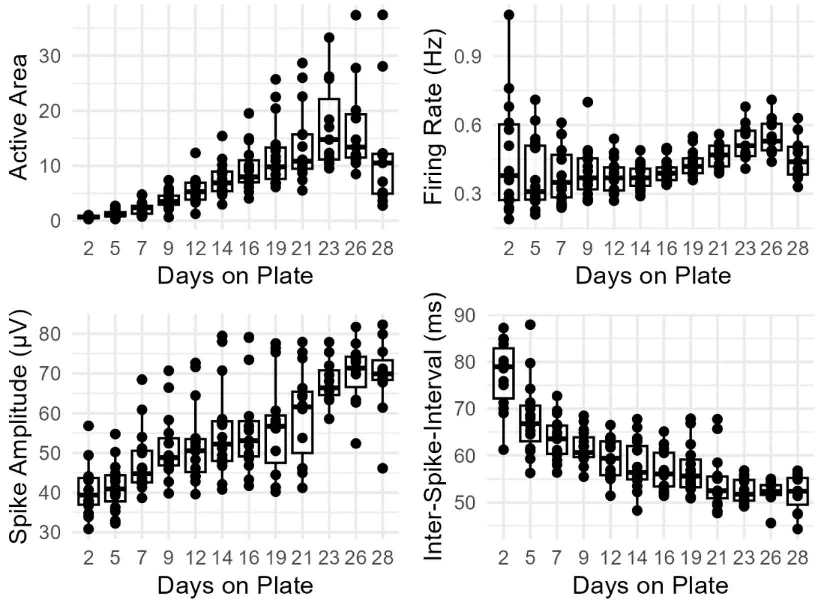


**Supplemental Figure 5. Ontogeny of general activity in control BrainSpheres over 28 days.** Graphs represent the number of days on plate across the x-axis for each of the different metrics describing general BrainSphere activity shown on the y-axis. The data represents an n of 11 BrainSpheres over a total of five hdMEA plates.

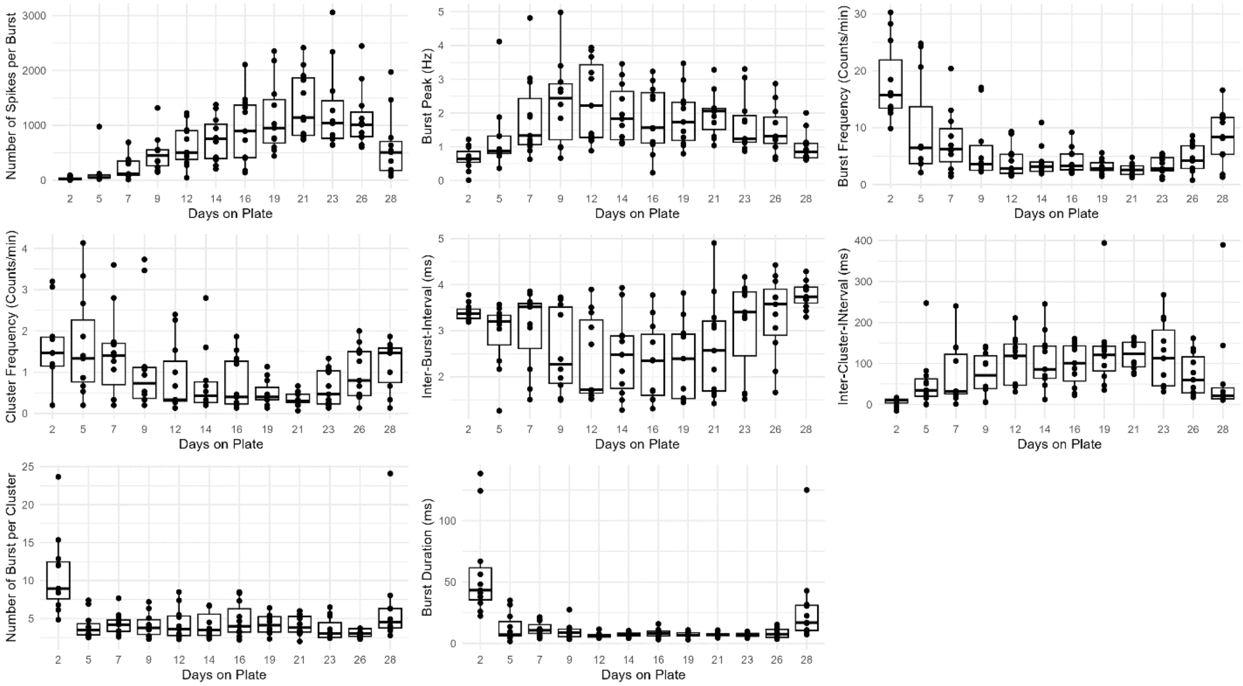


**Supplemental Figure 6. Ontogeny of Network activity in control BrainSpheres over 28 days.** Graphs represent the number of days on plate across the x-axis for each of the different metrics describing network burst-specific BrainSphere activity shown on the y-axis. The data represents an n of 11 BrainSpheres over a total of five hdMEA plates.

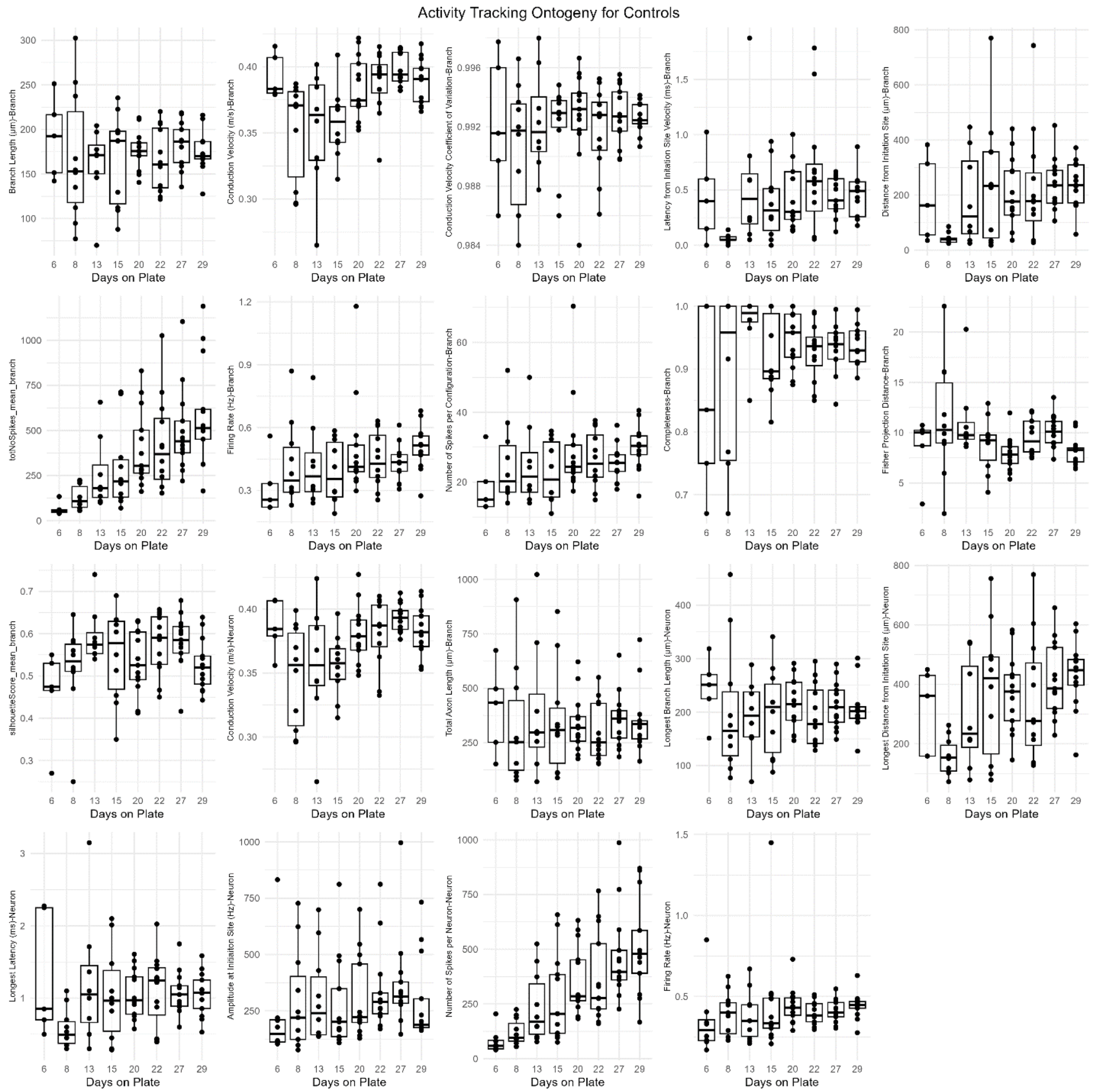


**Supplemental Figure 7. Ontogeny of axon potential propagation metrics in control BrainSpheres over 29 days.** Graphs represent the number of days on plate across the x-axis for each of the different metrics describing metrics utilized in axon tracking analysis in BrainSpheres shown on the y-axis. The data represents an n of 11 BrainSpheres over a total of five hdMEA plates.

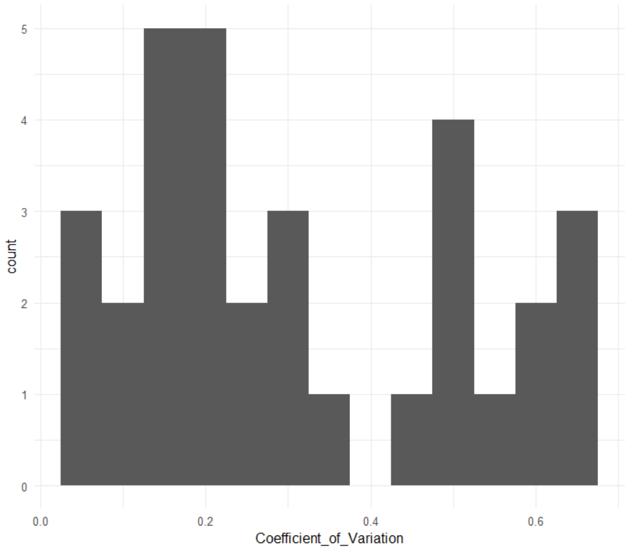


**Supplemental Figure 8. Distribution of the coefficient of variation values for all Endpoints.** The histogram represents the distribution of the values with bin widths equal to 0.1.
