## Supplementary figures and images for "Development of a network formation assay for developmental neurotoxicity hazard screening using 3D human iPSC derived BrainSpheres"

### Supplemental File 1

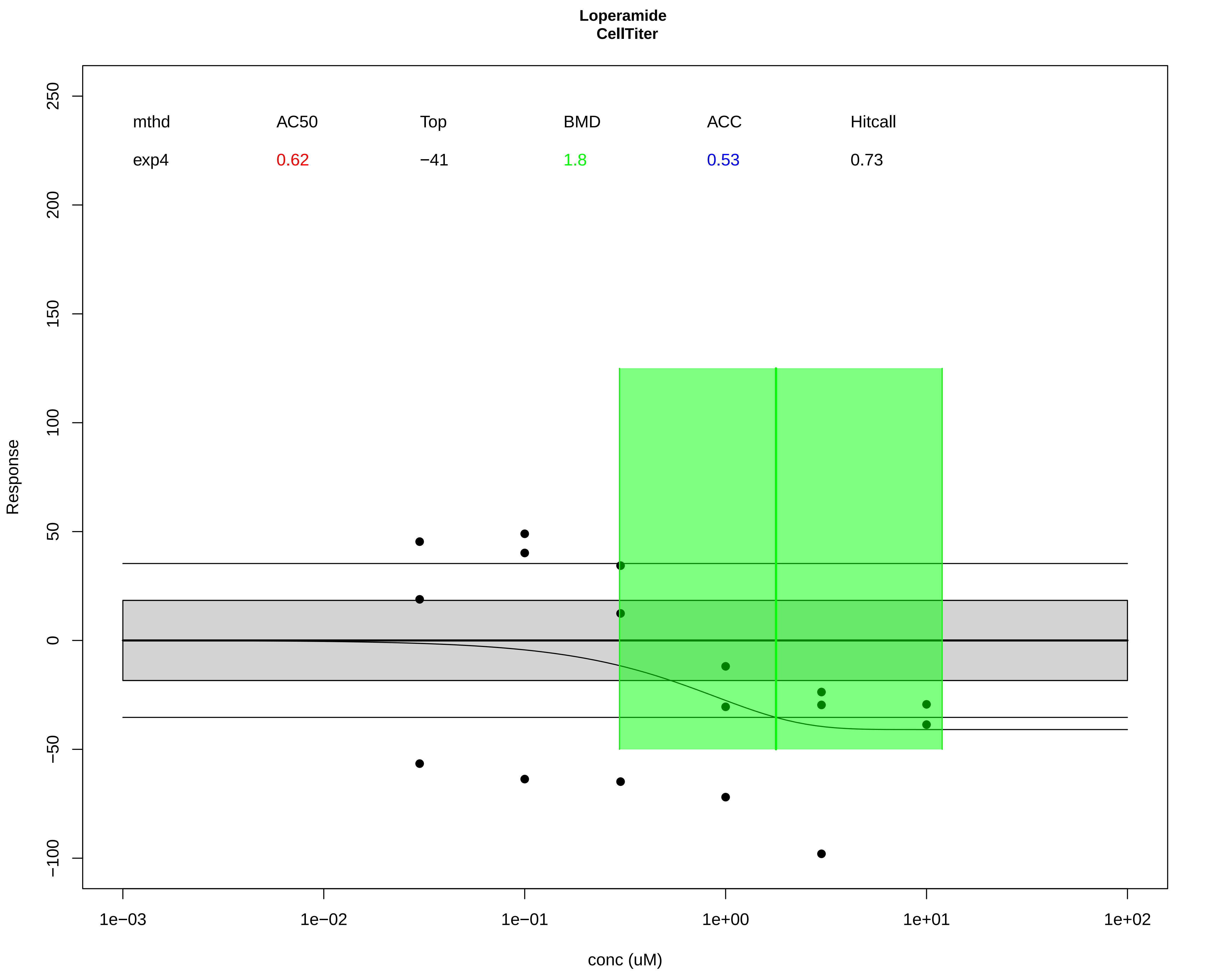

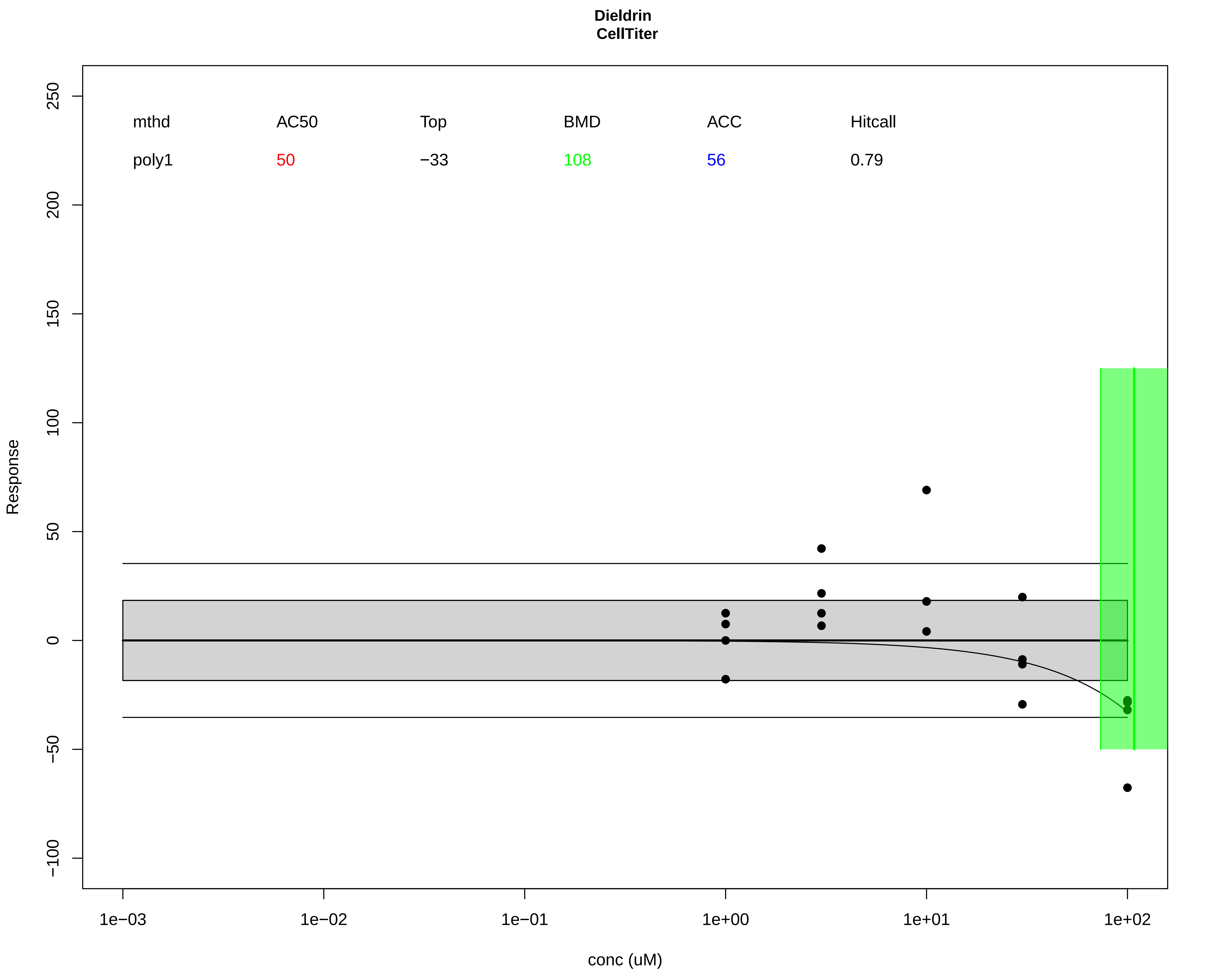

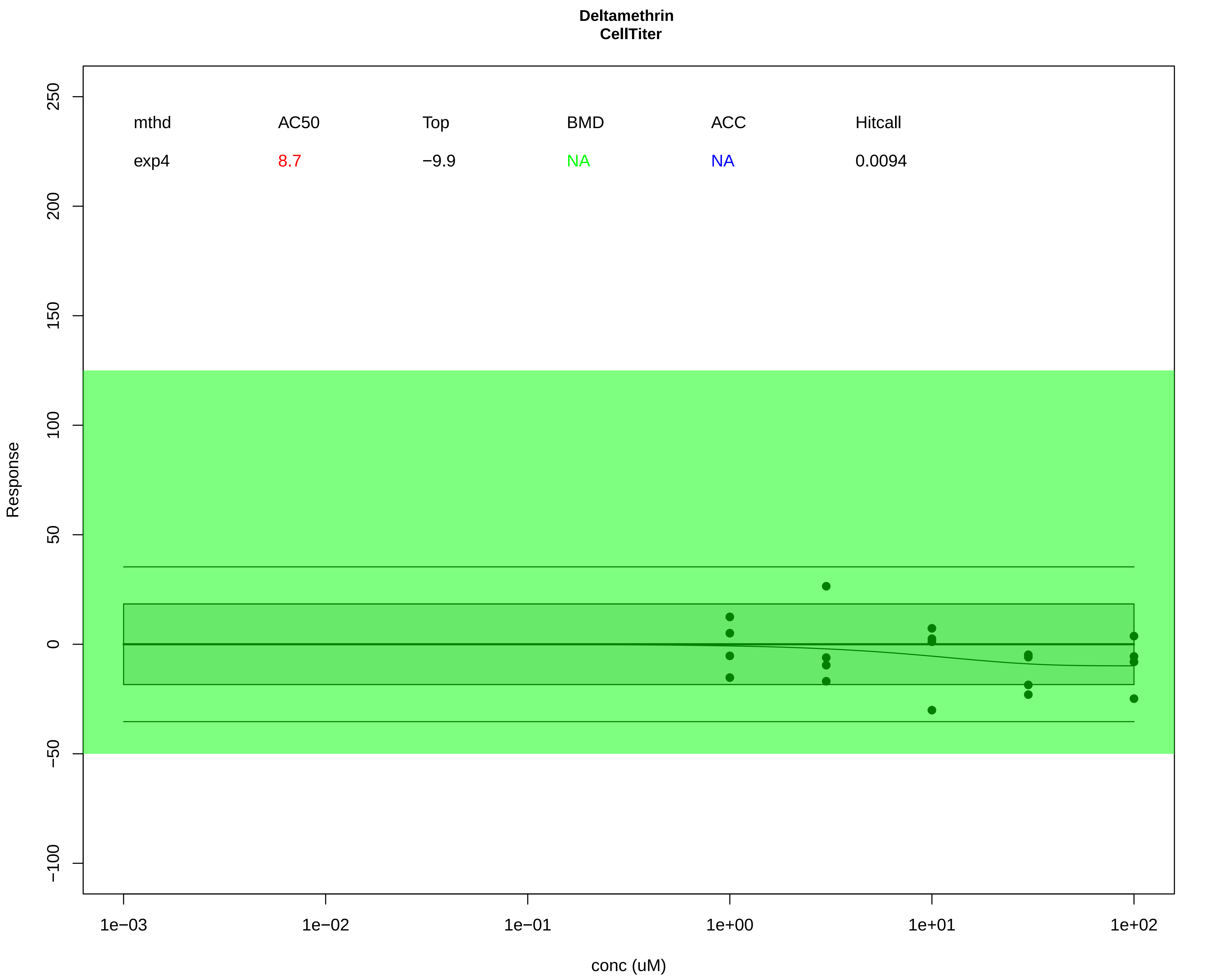

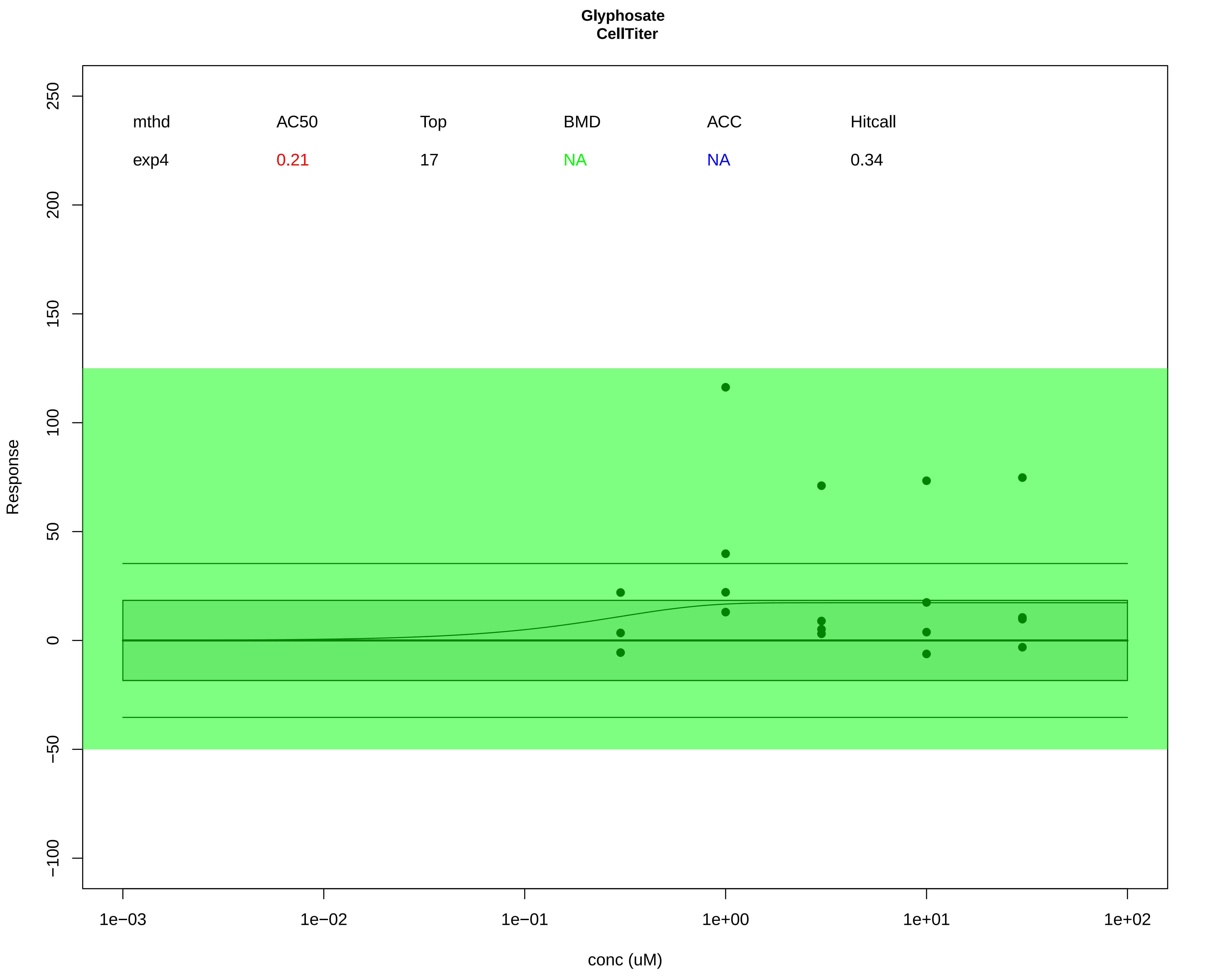

### Supplemental File 2

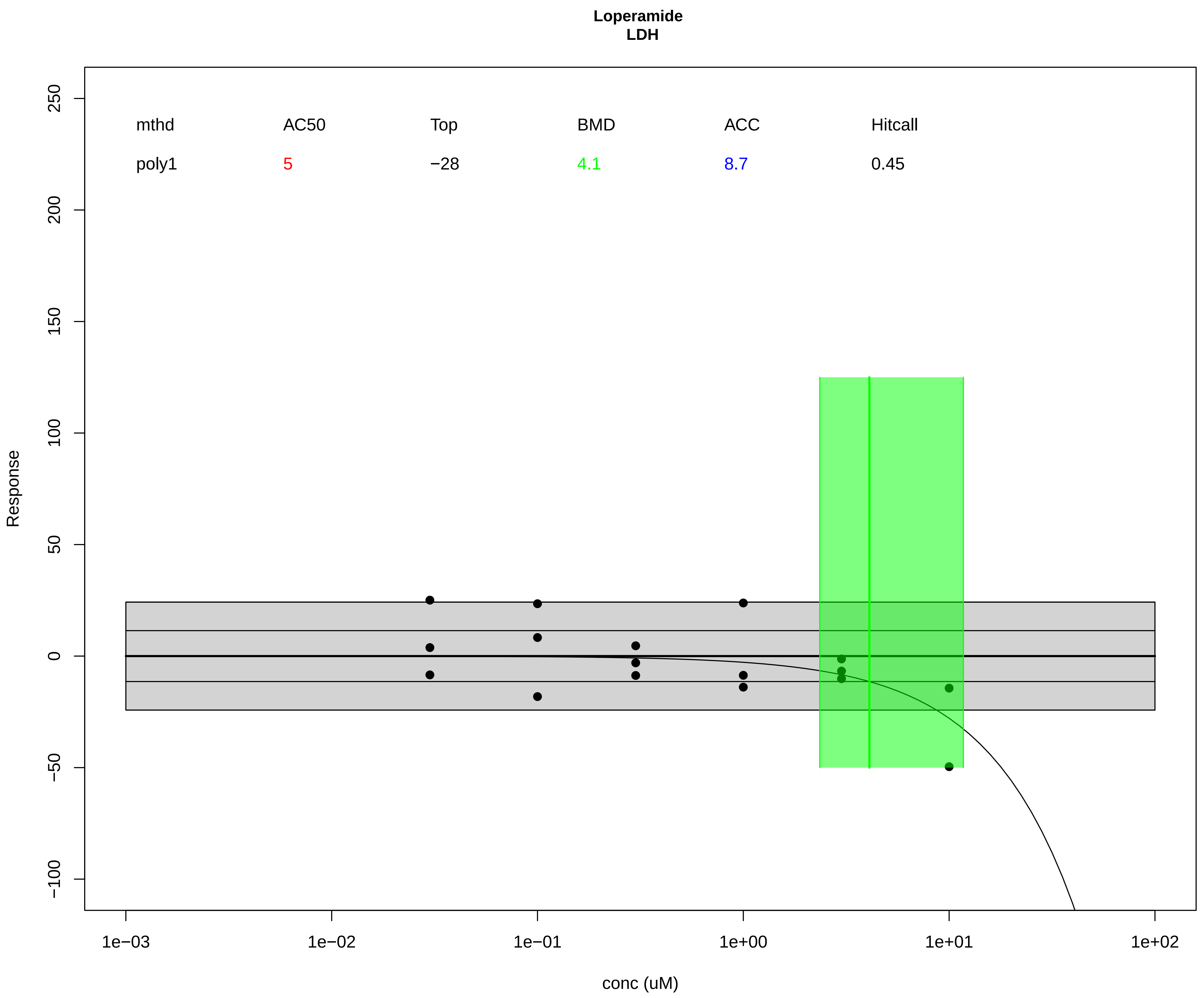

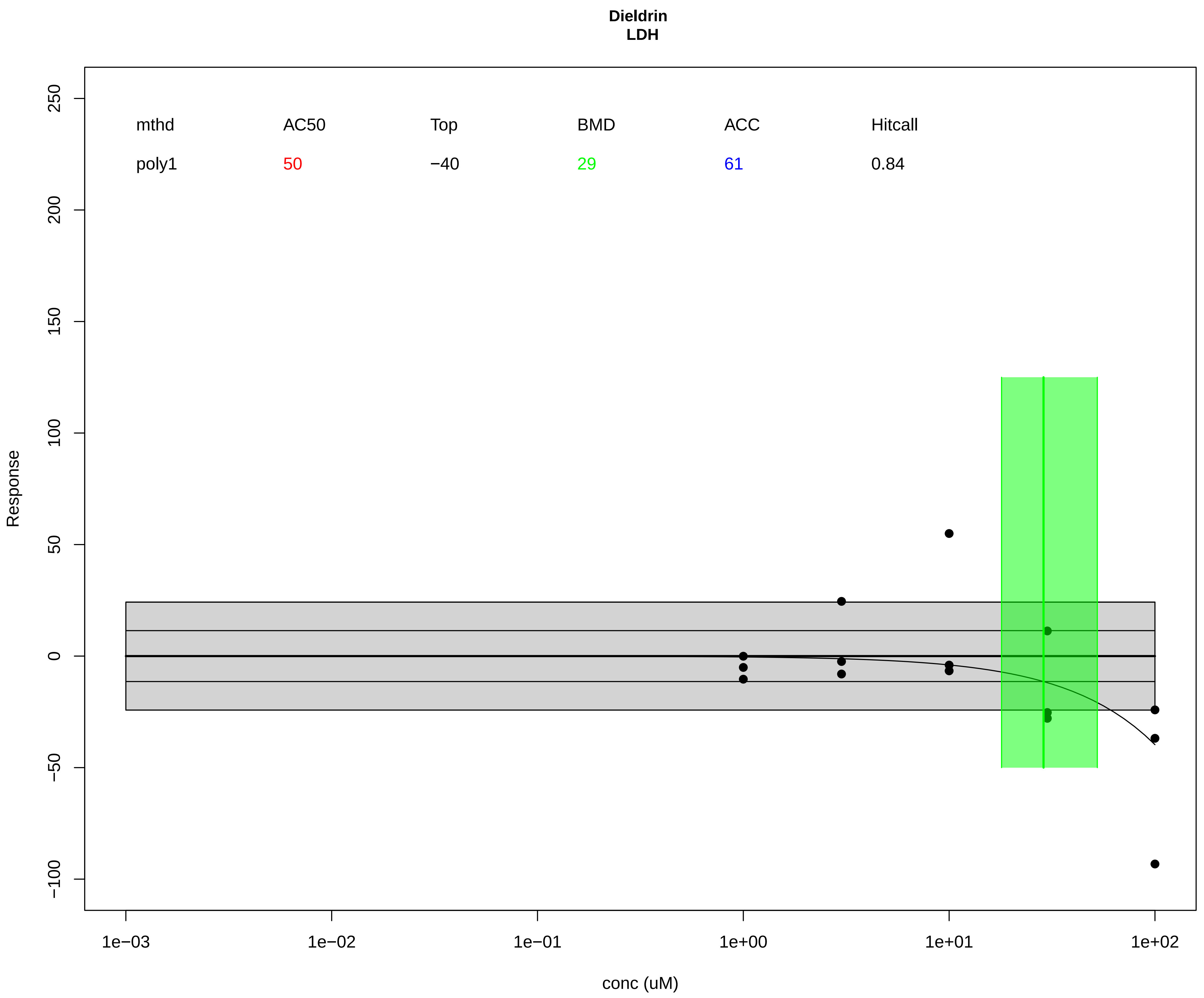

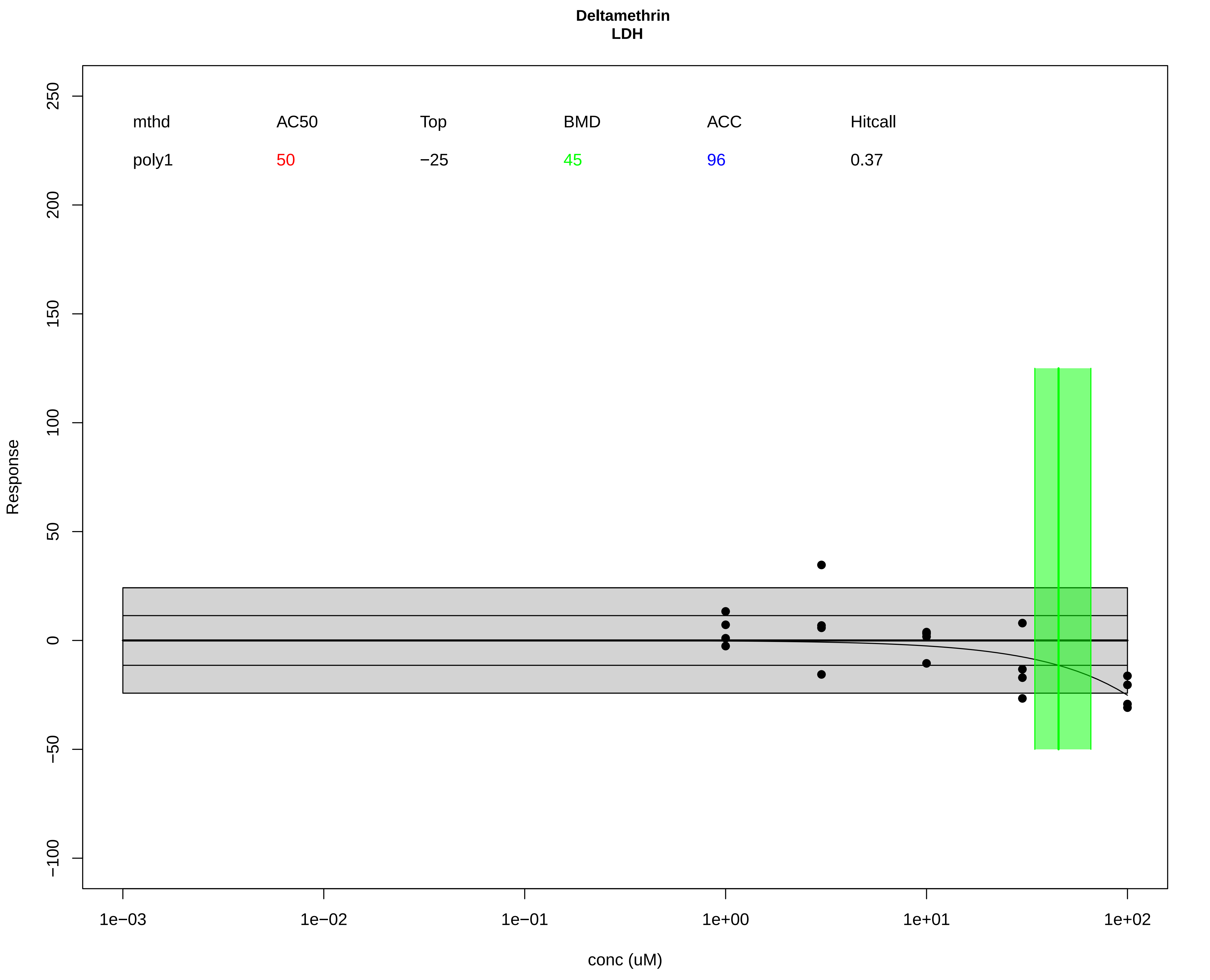

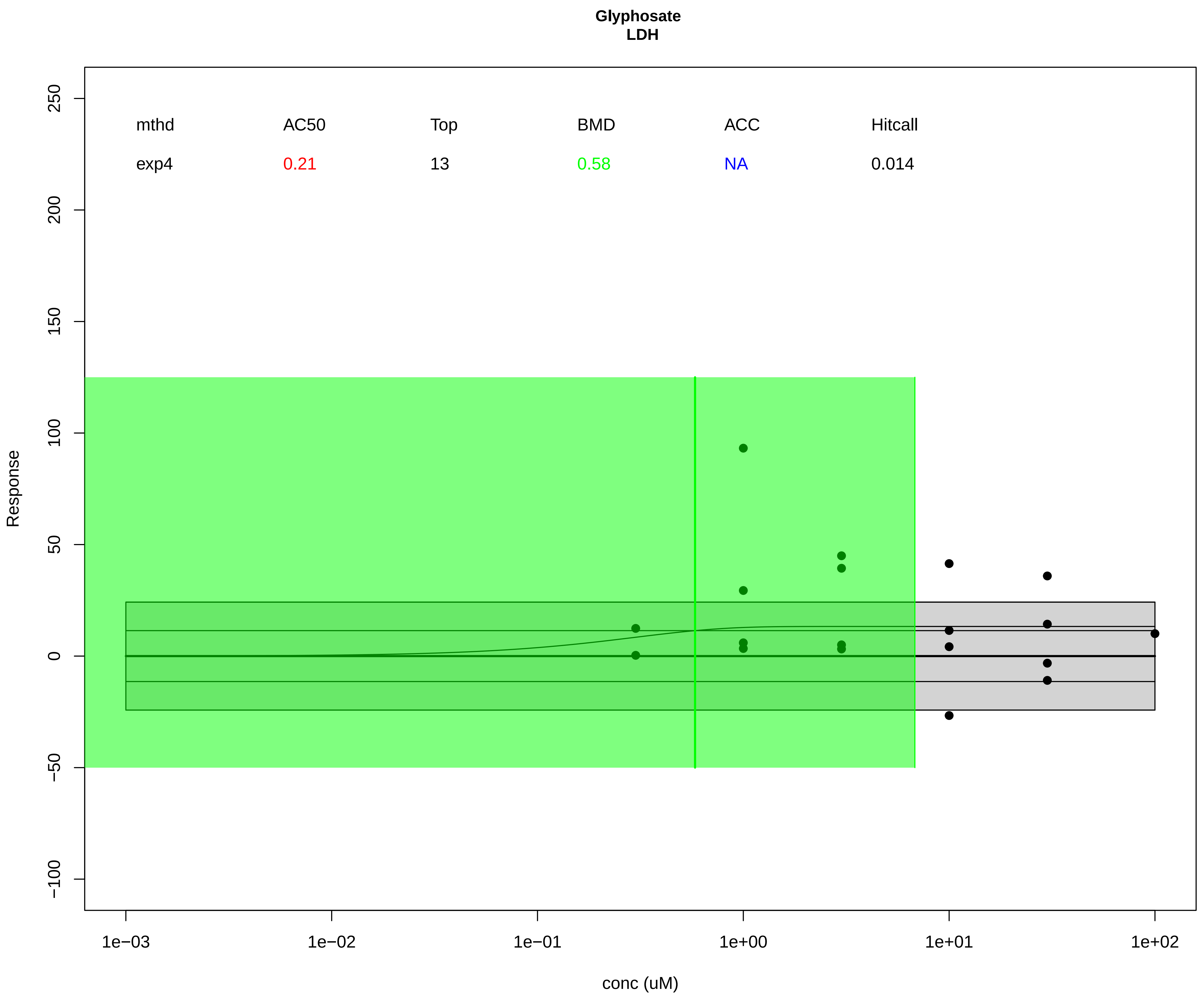

### Supplemental File 3

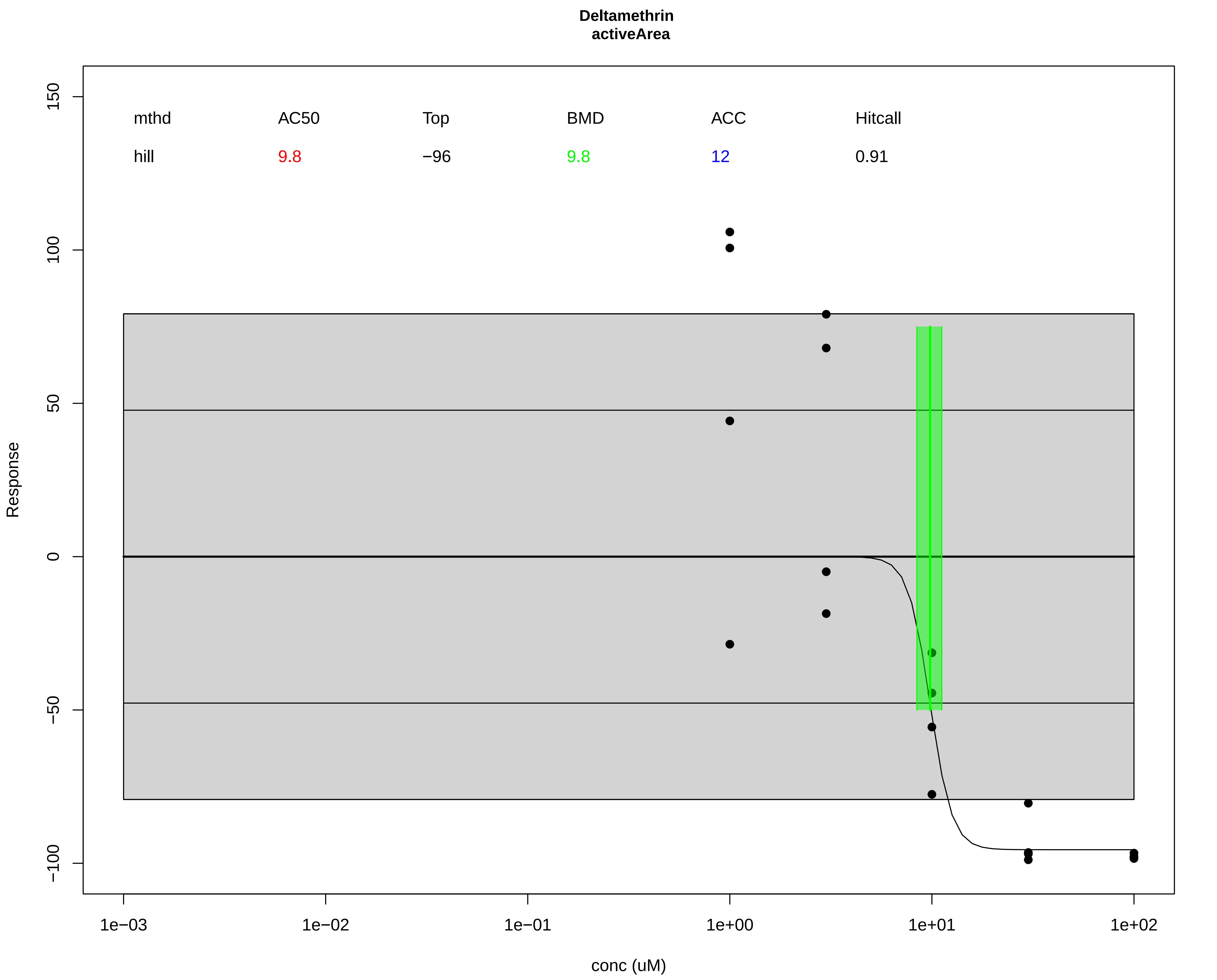

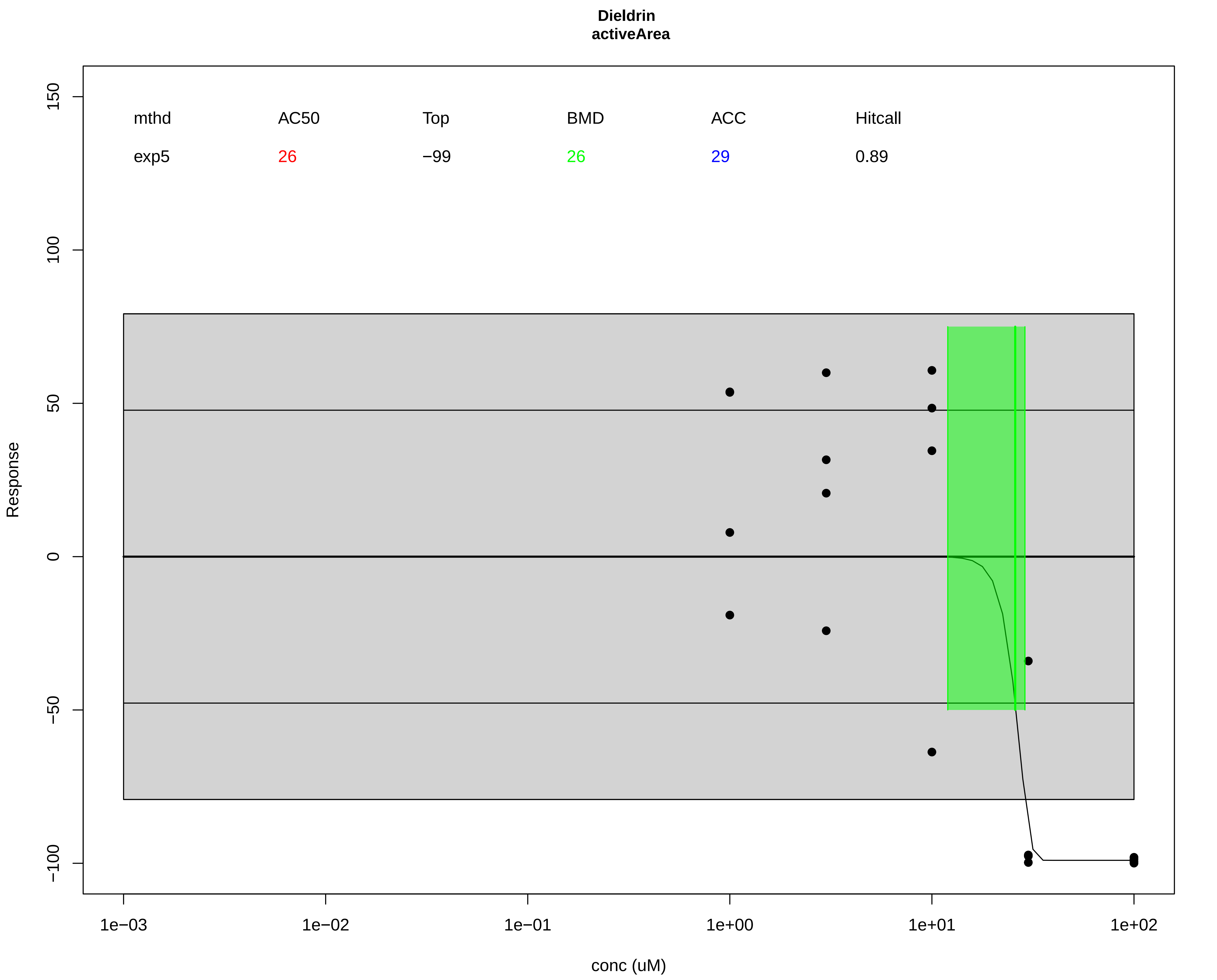

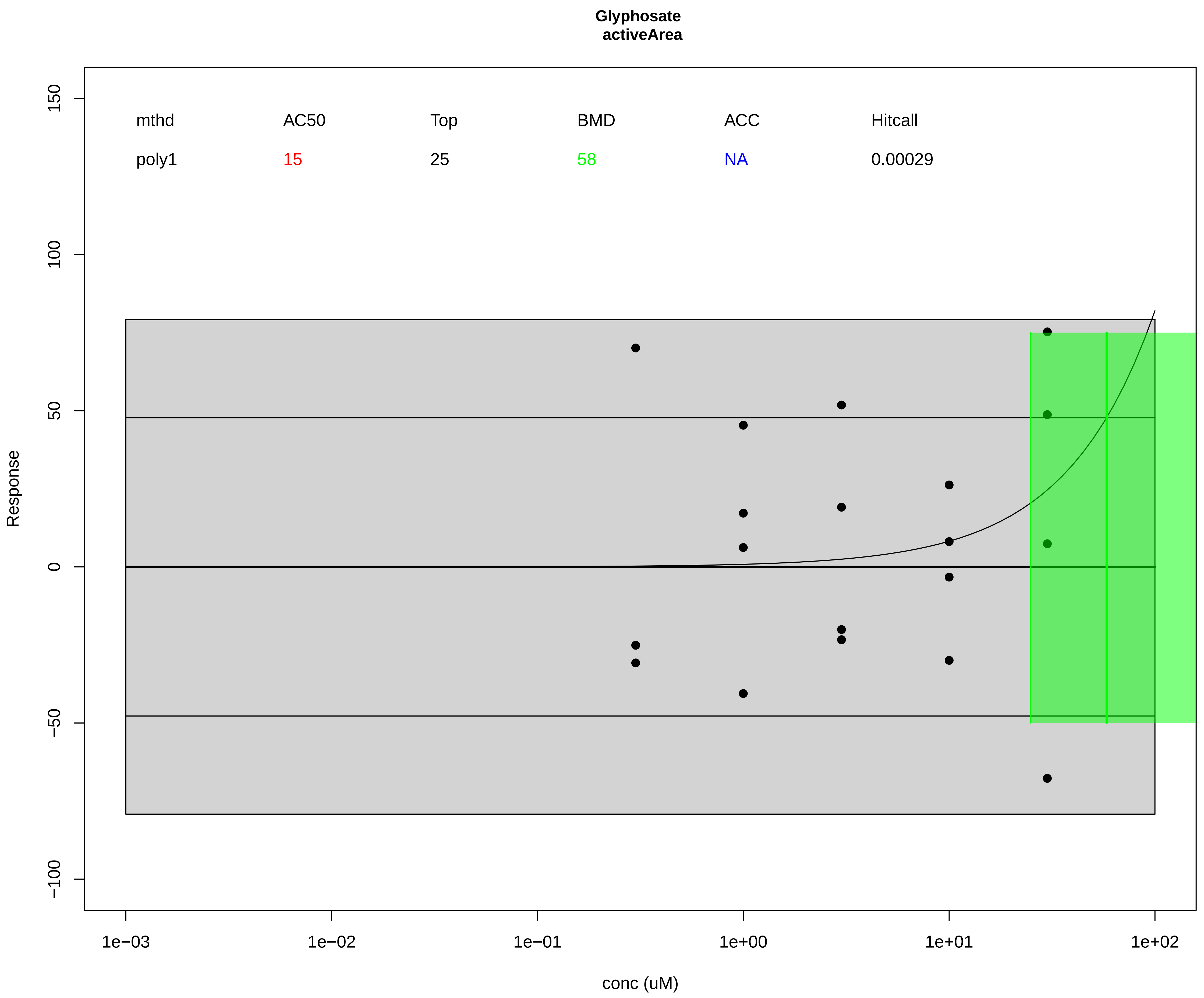

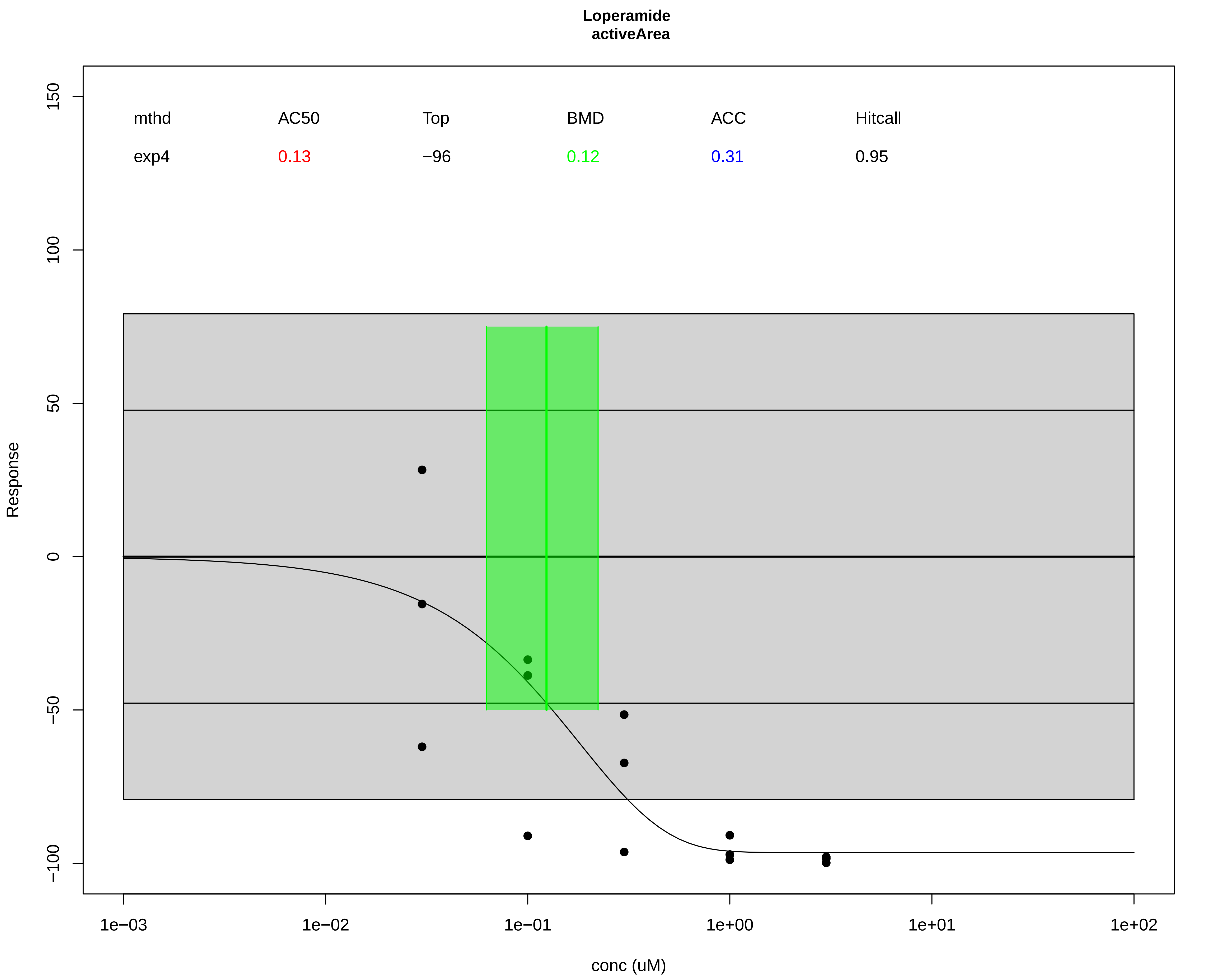

### Supplemental File 4

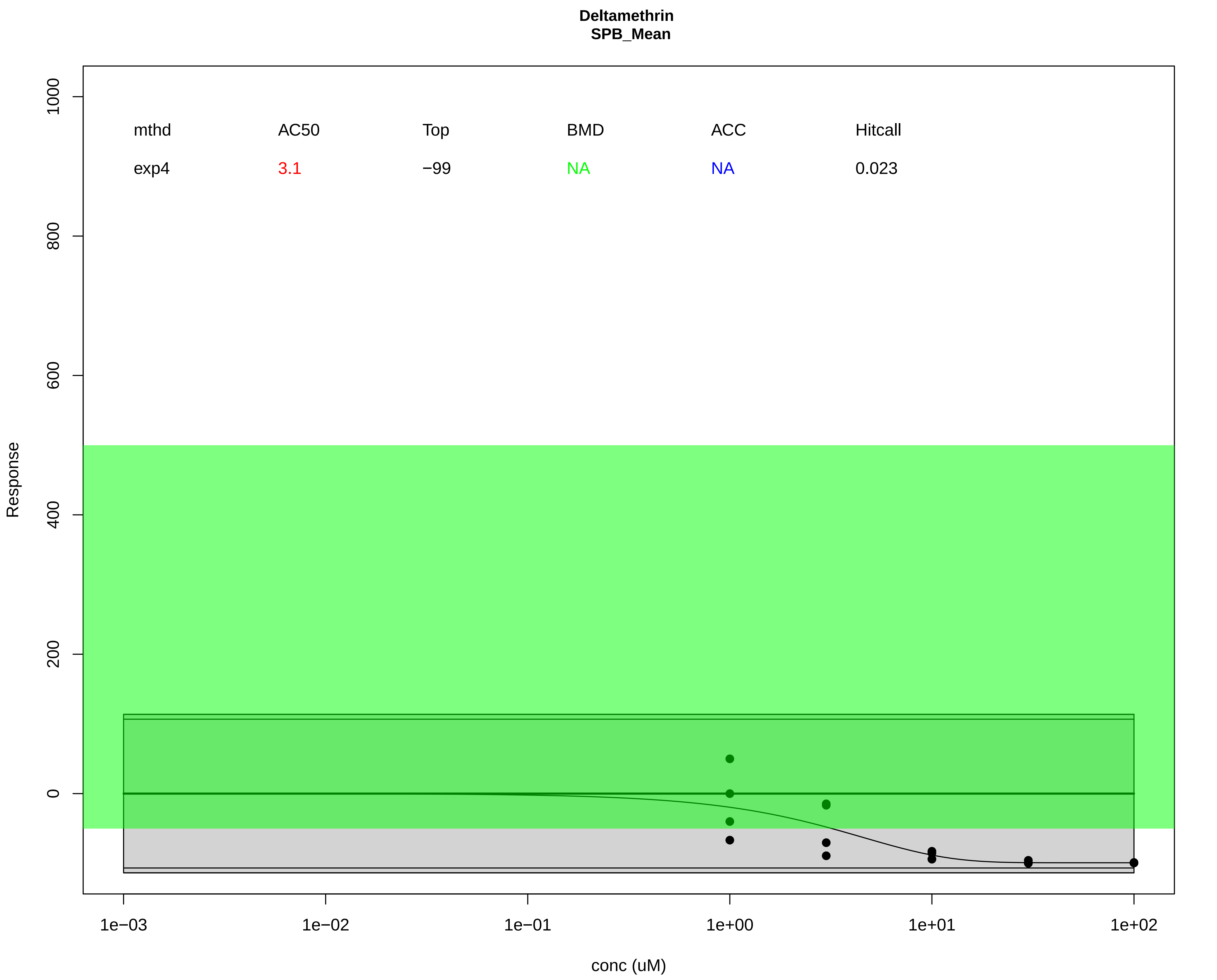

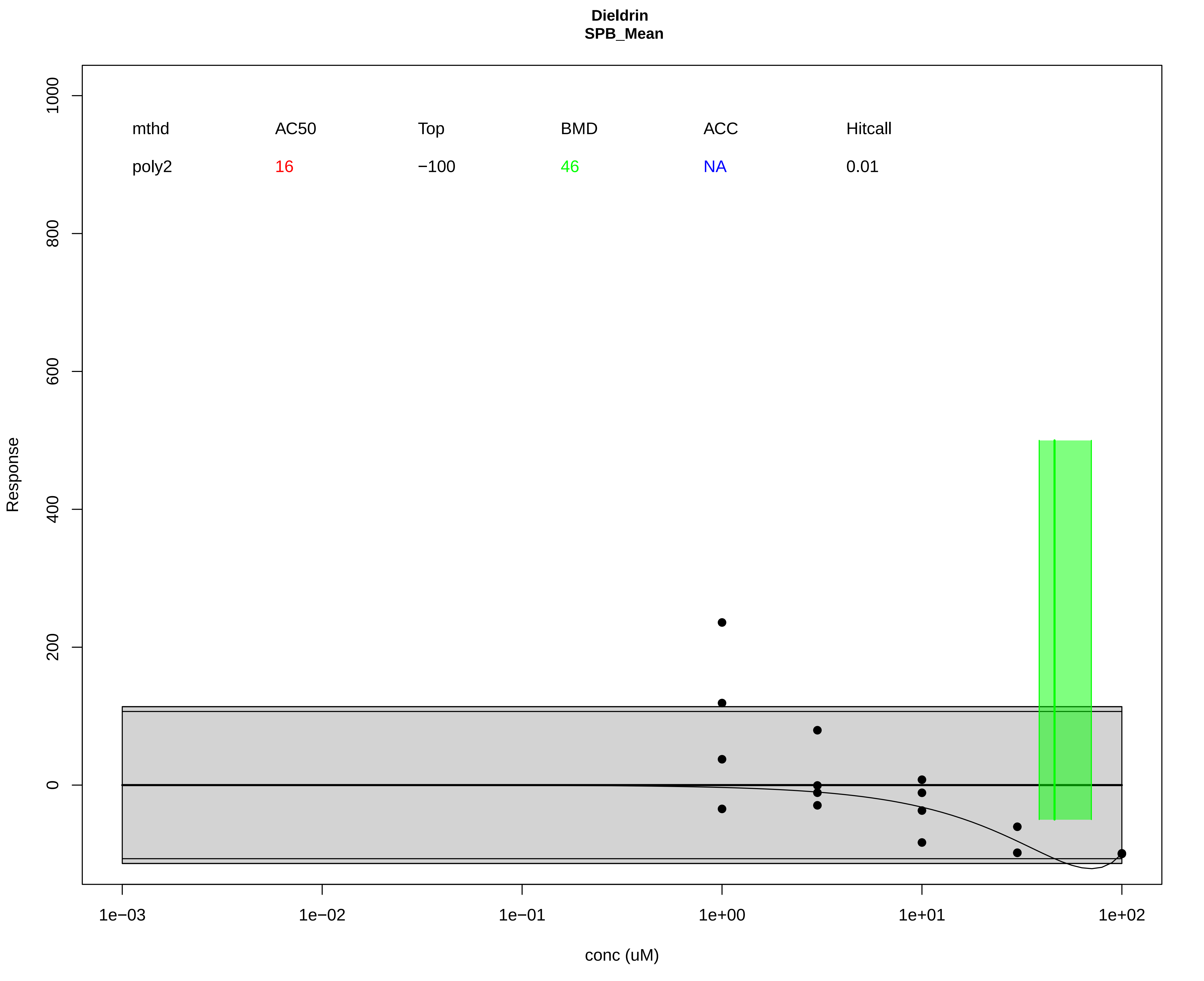

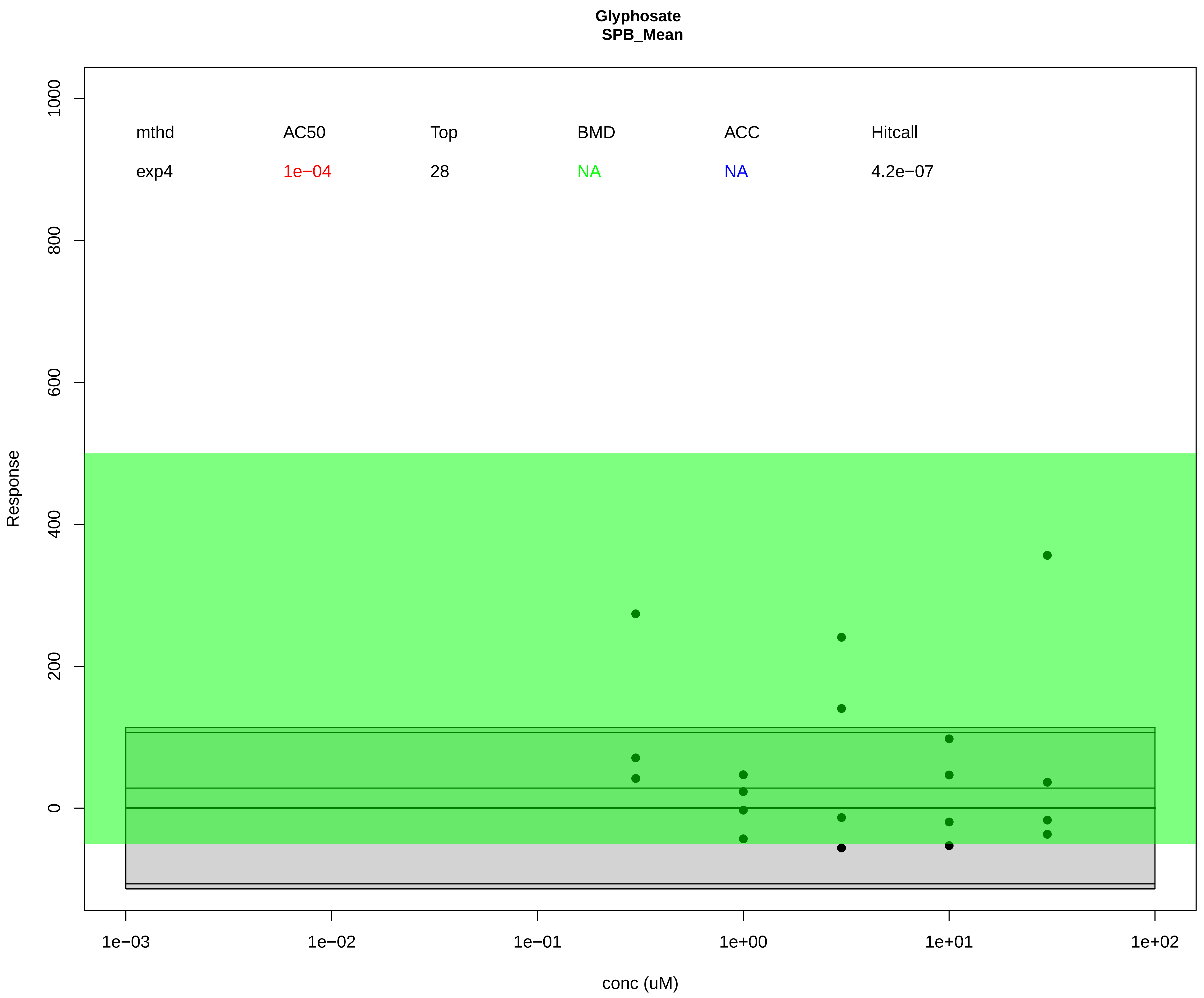

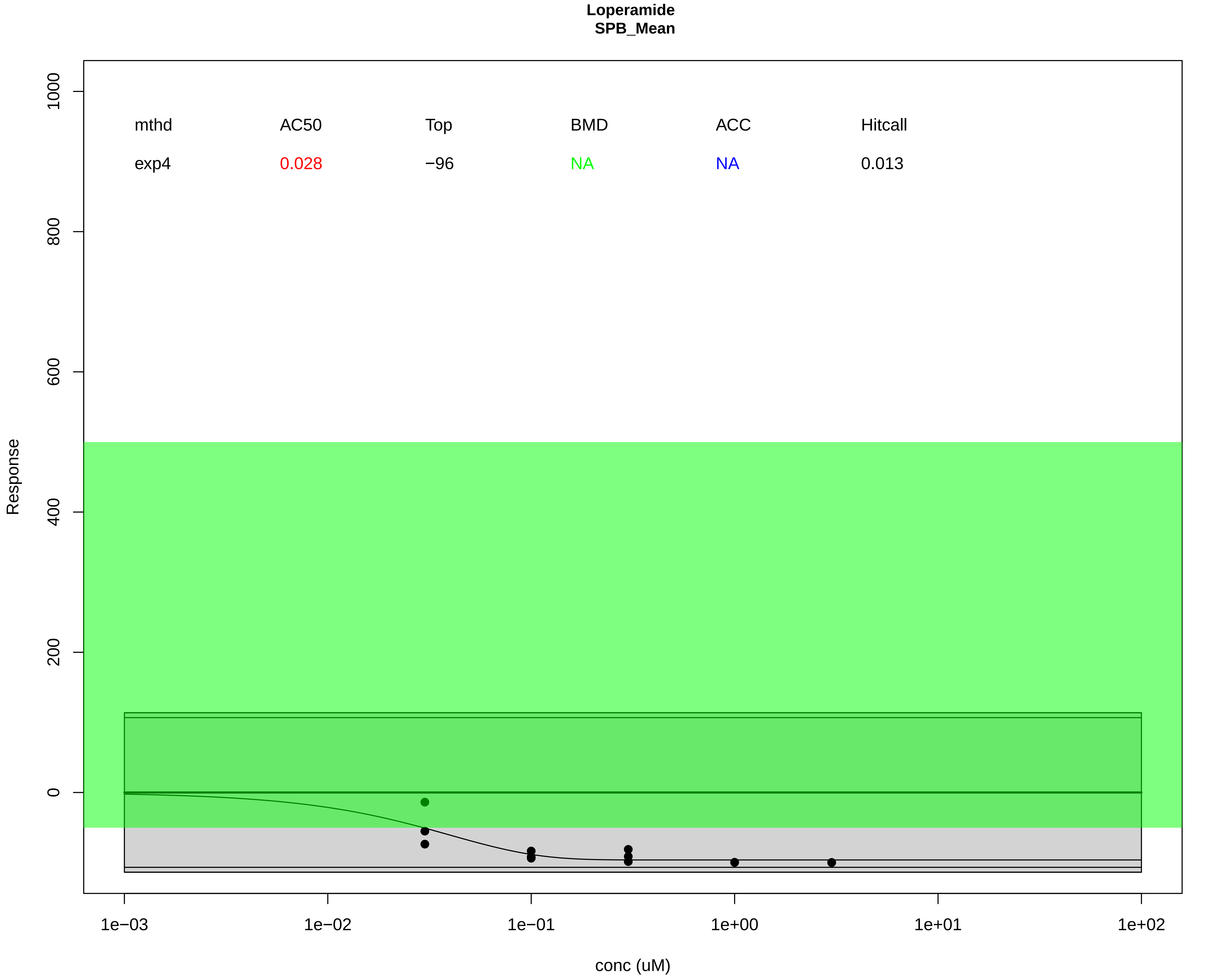

### Supplemental File 5

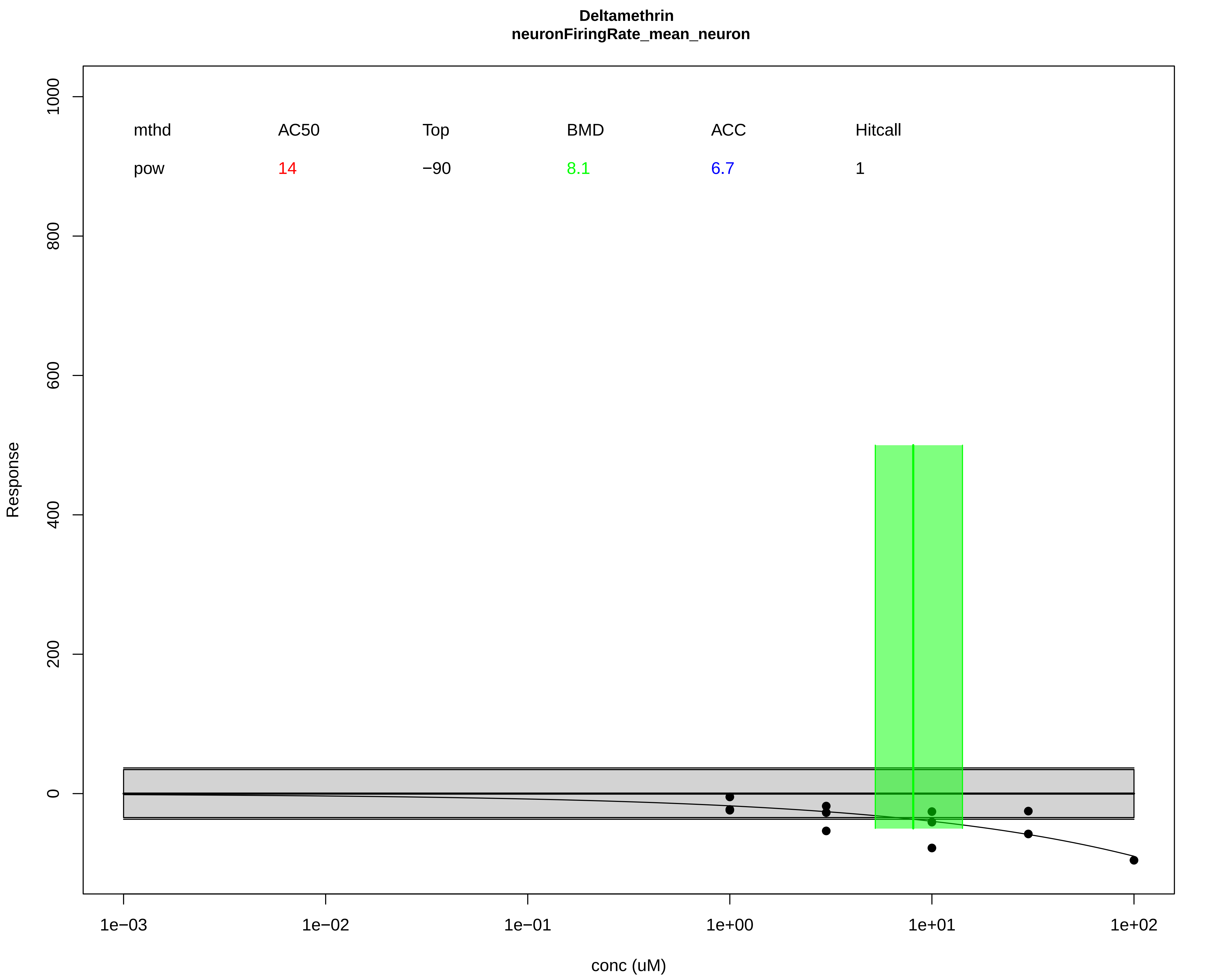

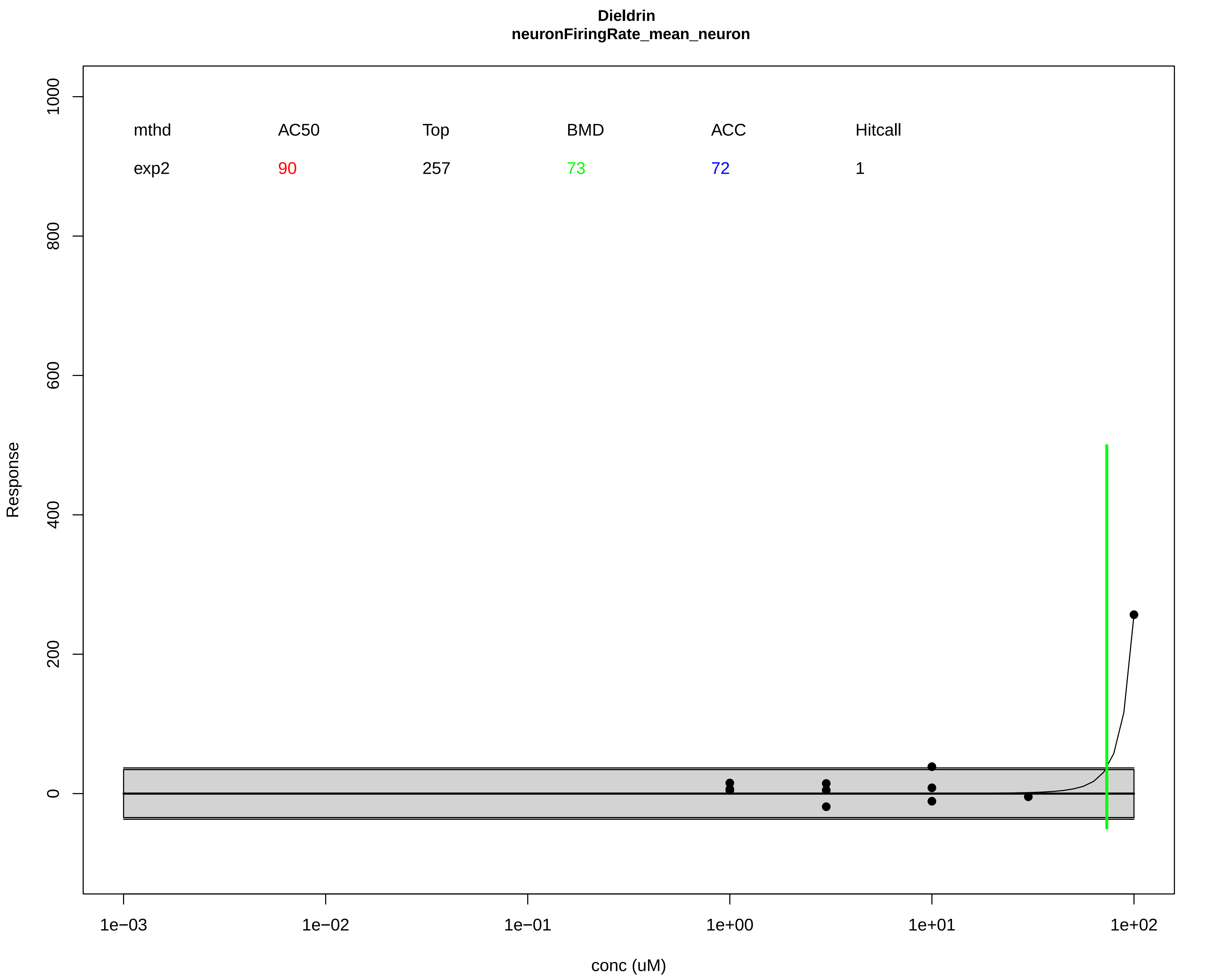

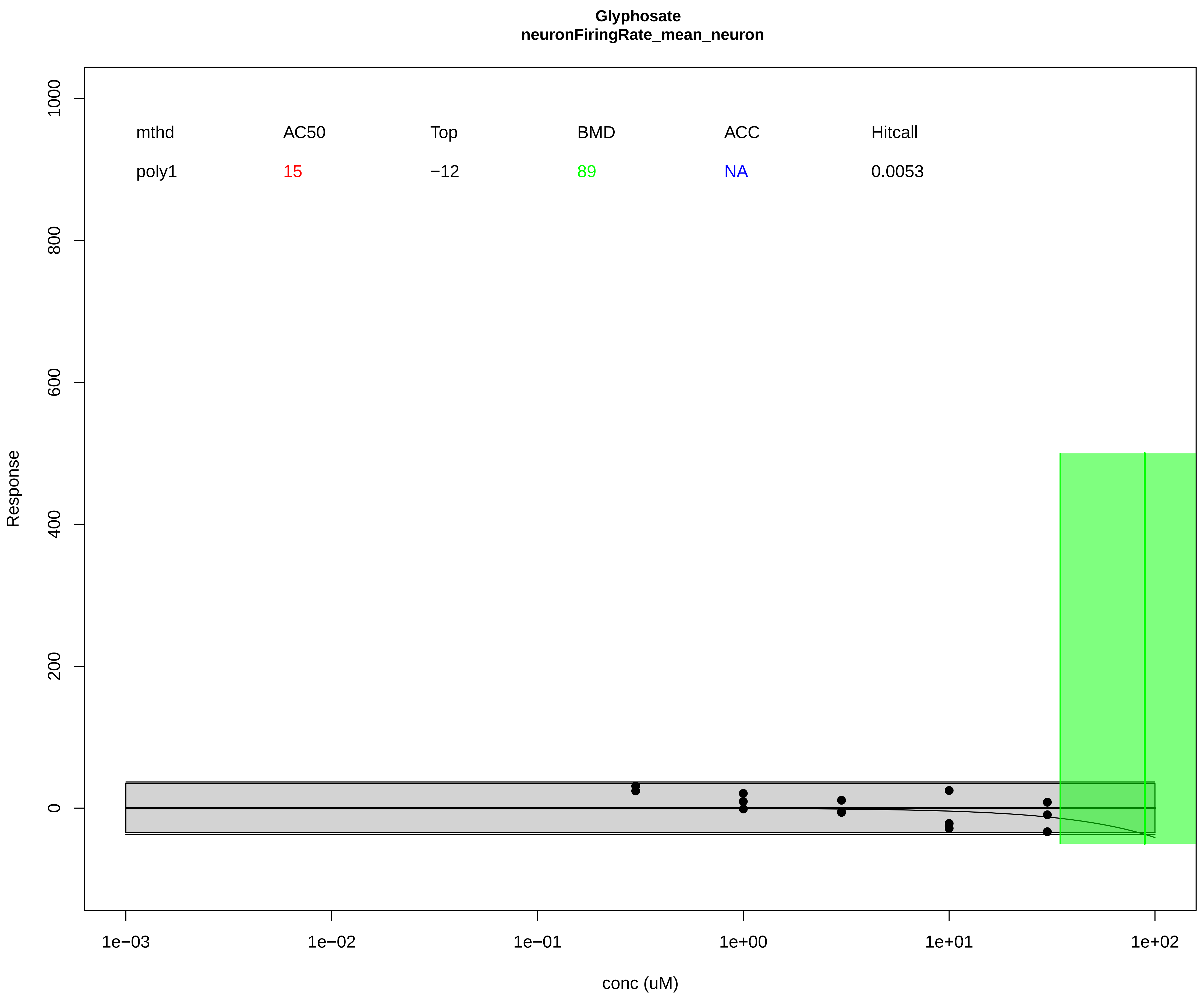

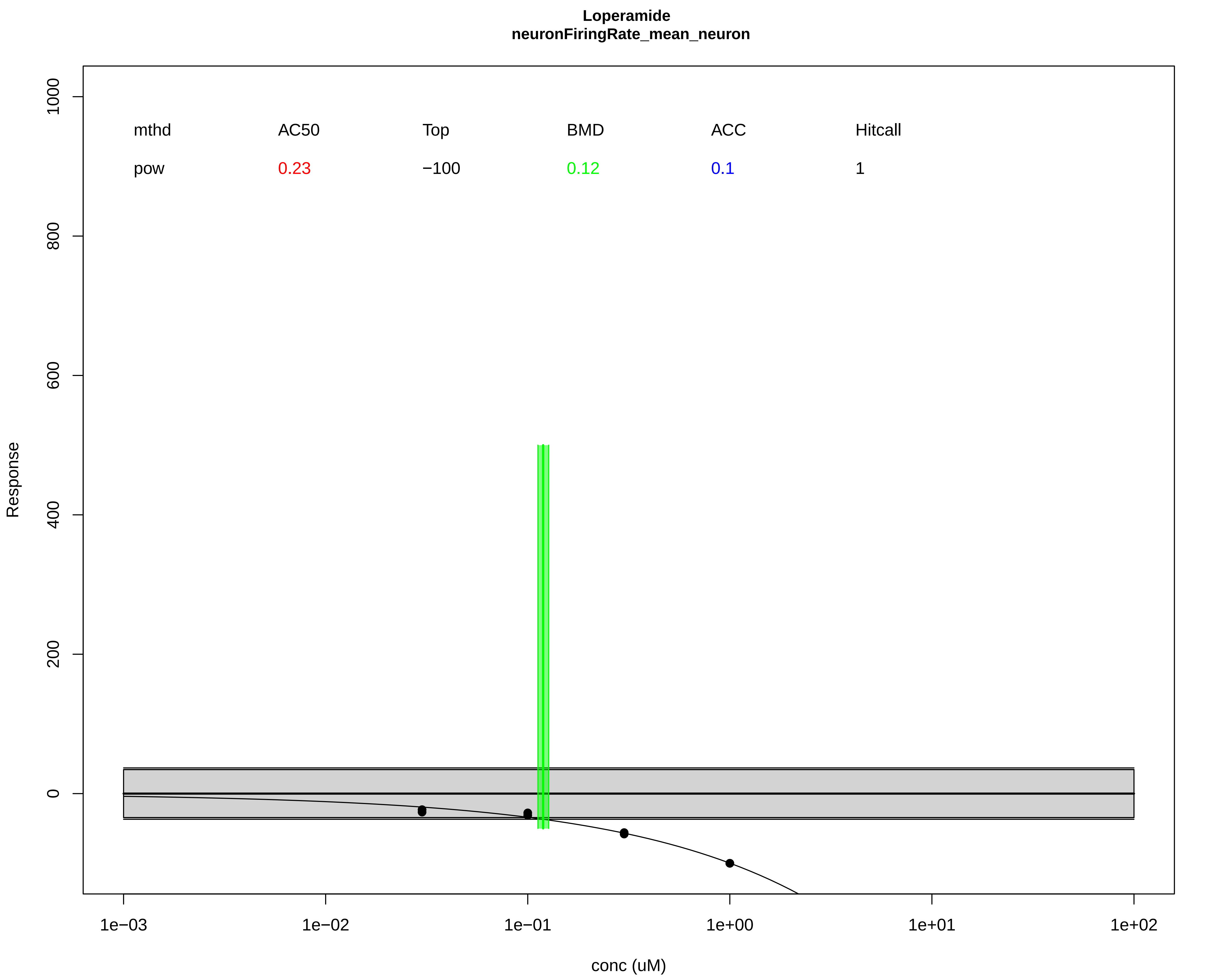
